# Glial-specific mitochondrial failure and redox imbalance drive regional vulnerability in Friedreich’s ataxia

**DOI:** 10.64898/2026.05.01.722124

**Authors:** Arabela Sanz-Alcázar, Marta Portillo-Carrasquer, Israel Manjarres-Raza, Maria Pazos-Gil, Fabien Delaspre, Jordi Tamarit, Juan P Bolaños, Joaquim Ros, Elisa Cabiscol

## Abstract

Friedreich’s ataxia (FA) is a rare autosomal recessive neurodegenerative disorder caused by reduced expression of frataxin, a mitochondrial protein important for iron–sulfur cluster assembly and mitochondrial homeostasis. Although FA has traditionally been attributed to neuronal dysfunction, increasing evidence suggests that glial cells play a critical role in disease progression, although their contribution remains poorly defined. Using the FXN^I151F^ mouse model, we investigated cell-type–specific metabolic and redox alterations in neurons and glial populations from the cerebrum, cerebellum, and dorsal root ganglia (DRG). Neuronal and glial-enriched fractions were isolated by immunomagnetic separation and analyzed for mitochondrial function, iron metabolism and reactive oxygen species (ROS). The analyses identified the DRG as the most severely affected region, exhibiting early and pronounced mitochondrial respiratory deficits, increased ROS, mitochondrial iron accumulation, lipid peroxidation, and reduced levels of glutathione peroxidase 4 and nuclear factor erythroid 2–related factor 2 in both neuronal and non-neuronal cells. These results highlight the vulnerability of sensory neurons and their supporting satellite glial cells. In contrast, in the cerebrum and cerebellum, astrocytes displayed earlier and more severe alterations than neurons, including impaired respiratory chain efficiency, disrupted complex I-III supercomplex interaction, elevated ROS, and hallmarks of ferroptosis. Neuronal abnormalities emerged later, suggesting that glial dysfunction precedes -or drives- neuronal pathology within the central nervous system. Overall, these findings reveal pronounced region and cell-type-specific vulnerabilities in FA and support the importance of targeting glial mechanisms—particularly iron dysregulation, oxidative stress, and ferroptosis— as targets for potential therapeutic strategies.

## Introduction

Friedreich’s ataxia (FA) is a rare autosomal recessive neurodegenerative disorder caused by mutations in the frataxin gene (*FXN*) on chromosome 9 (9q13), which encodes the mitochondrial protein frataxin (1,2). The majority of patients harbour a homozygous GAA triplet-repeat expansion within intron 1 of *FXN*, resulting in reduced frataxin expression (1,3,4). Approximately 4% of affected individuals are compound heterozygotes, carrying a GAA expansion on one allele and a point mutation or deletion on the other (5). Clinically, FA typically displays progressive neurological symptoms arising from degeneration of sensory neurons in the dorsal root ganglia (DRG) and lesions in the dentate nucleus of the cerebellum, leading to ataxia (6,7). Additional manifestations include muscle weakness, scoliosis, diabetes mellitus, and hypertrophic cardiomyopathy, the latter being the leading cause of mortality (4,7,8). Although disease onset usually occurs during childhood or adolescence (6,9), late-onset forms have also been described (10). Frataxin localizes to mitochondria (11), where its deficiency disrupts mitochondrial function, resulting in iron accumulation and increased oxidative stress driven by excessive production of reactive oxygen species (ROS). Recent studies have shown that frataxin acts as an allosteric activator of iron–sulfur (Fe–S) cluster biosynthesis by accelerating persulfide transfer from the cysteine desulfurase NFS1 to the scaffold protein ISCU (12–14). Mitochondrial iron accumulation has been reported across multiple frataxin-deficient models (15–22); however, its precise contribution to disease progression remains unclear. Moreover, how mitochondrial and iron-related alterations differentially affect specific neural cell types and regions is still poorly understood. Although FA pathology has traditionally been attributed to neuronal dysfunction in both the central nervous system (CNS) and peripheral nervous system (PNS), growing evidence supports a critical contribution of glial cells to disease pathogenesis (23,24). Once considered primarily passive support cells, glia is now recognized as active regulators of neuronal and essential physiological processes (25–30). The nervous system comprises highly specialized cell types—including neurons, astrocytes, and satellite glial cells (SGCs)—that differ markedly in metabolic demands, mitochondrial function properties, and vulnerability to stress (31,32). This perspective underscores glial dysfunction as a potential driver of FA progression and highlights the importance of cell-type–specific analyses in this neurodegenerative disease.

In this context, the DRG are uniquely suited for investigating neuron–glia interactions in FA, owing for their distinctive anatomy and pathological involvement. They contain pseudounipolar neurons traditionally classified by function (33); however, transcriptomic evidence suggests a far greater degree of molecular heterogeneity than previously appreciated (34). Among the non-neuronal cells, SGCs are the predominant subtype, accounting for 60–66% of all glia (35,36). Unlike Schwann cells, which are primarily associated with axons, SGCs form multi-cellular sheaths that tightly envelop neuronal somata, generating distinct morphological and functional units (37,38). The SGC-to-neuron ratio scales proportionally with neuronal volume, ranging from approximately 4:1 in small neurons to over 10:1 in the largest mammalian neurons (39,40). Proprioceptive neurons are particularly metabolically demanding (41,42) and rely predominantly on mitochondrial oxidative phosphorylation (OXPHOS), rendering them susceptible to dysfunction in FA (23,43,44). Unlike the CNS, the DRG lack a classical blood–brain barrier and are instead protected by a more permissive blood–DRG barrier (45). SGCs form an intimate “neuron–glial unit” by tightly enveloping neuronal soma, enabling bidirectional metabolic and signalling interactions (46). They regulate potassium homeostasis via Kir4.1 channels and supply metabolic substrates, like lactate and glutamine, to neurons (47,48). Furthermore, SGCs secrete pro-inflammatory cytokines such as IL-1β, IL-6, and TNF-α (49,50). In FA, DRG exhibit profound alterations, including the loss of large proprioceptive neurons and pronounced glial hypertrophy, hyperplasia, and mitochondrial biogenesis (7,23,28,43). Finally, dysregulation of iron-regulatory proteins, including ferritin and ferroportin, in both neuronal and glial cells further implicates the role of iron disrupted metabolism in disease progression (28,44).

Beyond the PNS, FA profoundly affects the CNS, particularly the cerebellum (51,52). While pathology is classically defined by dentate nucleus and Purkinje cell degeneration (51,53), granule neurons—the most abundant neuronal population in the brain—also contribute to cerebellar circuitry and metabolic demand (54,55). These neurons are supported by Bergmann glia (BG), specialized astrocytes that regulate glutamate homeostasis and provide redox protection (56,57). BG dysfunction is increasingly linked to cerebellar ataxia due to a failure to protect Purkinje neurons from oxidative stress (56). Similarly, cerebral pyramidal neurons, such as Betz cells, depend on protoplasmic astrocytes for metabolic and redox support (58–61). CNS neurons and astrocytes utilize distinct metabolic strategies. Neurons rely heavily on mitochondrial OXPHOS for ATP (62–64), while exhibited limited glycolytic capacity due to 6-phosphofructo-2-kinase/fructose-2, 6-bisphosphatase 3 (PFKFB3) repression and diversion of glucose to the pentose phosphate pathway for NADPH production (65–67). In contrast, astrocytes favor glycolysis, express high PFKFB3 levels, and supply lactate to neurons (59,61,68–71). Despite their reliance on OXPHOS, neurons generate less ROS than astrocytes due to more efficient coupling of complexes I and III (72,73). The efficiency of OXPHOS is influenced in part by the organization of electron transport chain complexes into supercomplexes. While complexes I, III, and IV remain catalytically active as free entities, their interaction into supercomplexes is proposed to enhance electron transfer efficiency and limit electron leakage (74,75). Astrocytes exhibit a higher proportion of free complex I relative to assembled supercomplexes than neurons (73), contributing to higher basal ROS production. Astrocytes compensate for this through robust antioxidant defenses, including glutathione (GSH) synthesis and elevated nuclear factor erythroid 2–related factor 2 (NRF2) expression (30,76,77), while also managing iron homeostasis (25,78,79). In FA models, frataxin-deficient astrocytes display increased superoxide production, lipid accumulation, and inflammatory cytokine secretion (80–83). Early activation of astrocytes and microglia in the cerebellum of YG8-800 mice suggests glial dysfunction arises at initial disease stages (80). Notably, interventions like insulin-like growth factor-1 can restore frataxin expression in astrocytes, ameliorating pathology and extending survival (84).

To investigate region and cell-type–specific mechanisms *in vivo*, we used the FXN^I151F^ mouse model. This mouse carries the I151F point mutation, equivalent to the human I154F pathological mutation and exhibits widespread frataxin depletion across tissues. FXN^I151F^ mice display systemic mitochondrial dysfunction, including impaired OXPHOS activity, disrupted redox and iron homeostasis, and progressive neurobehavioral deficits. Pronounced bioenergetic and oxidative stress abnormalities are observed in the DRG and cerebellum, with lesser—but still detectable—alterations in the cerebrum (20–22,85). These features make FXN^I151F^ mice a robust model for dissecting differential neuronal and glial responses to frataxin deficiency across vulnerable regions of the CNS and PNS. Although frataxin’s role in mitochondrial function is well established, its effects on redox balance, iron handling, and ferroptotic signalling in distinct neural cell populations remain incompletely understood. In this study, neurons and glial cells from the cerebrum, cerebellum, and DRG were isolated at two ages: 21 weeks, prior to the onset of observable neurological symptoms, and 39 weeks, when neurological deficits are clearly evident. With this approach we have been able to integrate analyses of bioenergetics, respiratory chain organization, oxidative stress, iron homeostasis, and the NRF2 pathway, with the goal of defining cell-specific pathophysiological mechanisms and identifying potential therapeutic targets.

## Materials and methods

### Animals

Animal experiments were performed following the National Guidelines (Generalitat de Catalunya and Government of Spain, article 33.a 214/1997), which adhere to the ARRIVE guidelines. The Experimental Animal Ethical Committee of the University of Lleida (CEEA) reviewed and approved the experimental protocols. All procedures took place at the animal facility, and euthanasia followed AVMA guidelines. FXN^I151F^ heterozygous mice (C57BL/6J-Fxnem10(T146T,I151F)Lutzy/J) were acquired from Jackson Laboratory (Bar Harbor, ME, USA, stock no. 31922), as previously reported (20). Homozygous WT and FXN^I151F^ mice were bred from heterozygous parents. Animals were housed in standard ventilated cages with a 12 h light/dark cycle and free access to standard chow. Weekly weight measurements were taken. Genotyping was done by sequencing PCR products from tail DNA, as previously described (20).

### Immunomagnetic purification of neurons and astrocytes from cerebrum and cerebellum

The protocol for preparing single-cell suspensions from murine cerebellum or cerebrum was adapted from (73,86). Briefly, dissected cerebellar or cerebral tissue was rinsed in Solution A (Earle’s Balanced Salt Solution (EBSS); Fisher, ref. 11540616) supplemented with 0.4% bovine serum albumin (BSA, Sigma, ref. A9647) and 0.026 mg/mL DNAse I grade II (Roche, ref. 104159)) to remove excess blood. The tissue was then longitudinally sliced, further divided, and transferred to a conical tube containing Solution B (EBSS supplemented with 0.028% BSA, 0.052 mg/mL DNAse I grade II, and 1 mg/mL Trypsin; Sigma, ref. T4799). Enzymatic digestion was performed by incubation in a 37°C water bath with gentle agitation for 5 min, followed by mechanical dissociation through repeated pipetting. The cell suspension was then incubated for an additional 10 min at 37°C with agitation. Trypsin activity was subsequently quenched by the addition of fetal bovine serum to a final concentration of 10%. Following centrifugation at 700*×g* for 5 min at 4°C, the supernatant was discarded, and the cell pellet was resuspended in Solution A through gentle pipetting with a siliconized Pasteur pipette. After allowing larger debris to settle, the upper cell suspension was collected and filtered through a 70 μm cell strainer (ClearLine, ref. 141379C). This sedimentation and filtration procedure was repeated twice to maximize cell yield and remove residual debris. The filtered cell suspension was centrifuged at 700 *× g* for 3 min, the supernatant was aspirated, and the pellet was resuspended in a red blood cell lysis solution (Miltenyi Biotec, ref. 130-094-183) to remove erythrocytes. The resulting cell suspension was centrifuged at 500 *× g* for 10 min, and the final pellet was resuspended in Dulbecco’s phosphate-buffered saline (DPBS) (Gibco, ref. 14190-136) with 0.5% BSA, yielding a homogeneous single-cell suspension. Specific cell populations were then isolated from this suspension using MACS technology (Miltenyi Biotec) according to the manufacturer’s instructions. Neurons were isolated using the Neuron Isolation Kit Mouse (Miltenyi Biotec, ref. 130–115–390), and astrocytes were selected using the Anti–ACSA–2 MicroBead Kit, Mouse (Miltenyi Biotec, ref. 130–097–678). A schematic representation is shown in Supplementary Fig. S1A.

### Immunomagnetic purification of neurons and non-neuronal cells from DRG

DRG were dissected from mice into a Petri dish containing GHEBS buffer (137 mM NaCl, 2.6 mM KCl, 25 mM glucose, 25 mM HEPES, 100 μg/mL penicillin/streptomycin), and adhering nerve tissue was meticulously removed as previously described (21). Approximately 30–40 ganglia were harvested per mouse. The dissected DRG were transferred to a conical tube and enzymatically digested with 1 mg/mL collagenase (Roche, ref. 10103578001) and 1 mg/mL dispase II (Sigma, ref. D4693), both dissolved in GHEBS buffer, for 60 min at 37°C in a water bath. Following digestion, the cells were washed and homogenized in DPBS supplemented with 0.5% BSA and 3 mg/mL DNAse I grade II. The resulting suspension was filtered through a 70 µm cell strainer (pluriSelect). The filtrate was centrifuged at 400 × *g* for 5 min, and the pelleted cells were resuspended in DPBS with 0.5% BSA and a red blood cell removal solution. After centrifugation at 500 *× g* for 10 min, the final pellet was resuspended in DPBS with 0.5% BSA, yielding a homogeneous single-cell suspension. Subsequently, specific cell populations were isolated from the homogeneous cell suspension using MACS Technology according to the manufacturer’s instructions. Neurons were selected using the neuron-specific Neuron Isolation Kit, and non-neuronal cells were obtained from the negative fraction of this separation. A schematic representation is shown in Supplementary Fig. S1B.

### Mitochondrial respiration in neurons and astrocytes

Oxygen consumption rate (OCR) and extracellular acidification rate (ECAR) were measured in neurons and astrocytes isolated from the cerebrum using the Seahorse XF analyzer (Agilent). Prior to the assay, neuronal and astrocytic cells obtained as described above, were quantified using the LUNA-II™ automated cell counter and subsequently seeded at a density of 2×10^6^ cells per well into XF24 cell culture microplates (Agilent, ref. 100777-004). These plates were pre-coated with poly-D-lysine (Sigma, ref. P1149), and cells were seeded in Dulbecco’s modified Eagle’s medium (DMEM, pH 7.4, Agilent, ref. 103575-100) supplemented with 2 mM glutamine (Agilent, ref. 103579-100), 1 mM pyruvate (Agilent, ref. 103578-100), and 10 mM glucose (Agilent, ref. 103577-100). The plates were then incubated at 37°C in a non-CO₂ incubator for 35–40 min to allow cell adherence. OCR and ECAR measurements were performed according to the manufacturer’s instructions provided with the Mito Stress Test Kit. Mitochondrial respiratory chain inhibitors were sequentially introduced into each well via the automated injection system: 1.5 µM oligomycin A (Sigma, ref. 75351) to inhibit ATP synthase (complex V), 1.5 µM (neurons) or 5 µM (astrocytes) carbonyl cyanide-p-trifluoromethoxyphenylhydrazone (FCCP, Sigma, ref. C2920) to uncouple the mitochondrial membrane potential, and 1 µM rotenone (Sigma, ref. R8875) and 1 µM antimycin A (Sigma, ref. A8674) to inhibit complex I and complex III, respectively. Data acquisition, analysis, and graphical representation were performed automatically using the Seahorse analytics software. For the normalization, cells were fixed in situ with 4% paraformaldehyde and subsequently stained with Hoechst dye to determine cell number using the Operetta CLS high-content microscope system (PerkinElmer).

### Mitochondrial isolation and Blue native gel electrophoresis (BNGE) of neurons and astrocytes

Mitochondrial isolation and BNGE was performed as described (73) with modifications. Isolated cerebral neuronal and astrocytic cell suspensions were centrifuged, and the resulting cell pellets were cryopreserved at −80 °C. For subsequent processing, frozen pellets were homogenized at 4°C using a teflon homogenizer in Buffer A containing 83 mM sucrose (Sigma, ref. 18219) and 10 mM MOPS (Sigma, ref. M-3183), pH 7.2. An equal volume of Buffer B (250 mM sucrose, 30 mM MOPS) was then added to the homogenate. This suspension was centrifuged at 1,000 *× g* for 5 min at 4°C to remove nuclei and residual intact cells. The supernatant was then subjected to centrifugation at 12,000 *× g* for 2 min at 4°C, leading to the sedimentation of the mitochondrial fraction. The mitochondrial pellet was resuspended in lysis buffer containing 1 M 6-aminohexanoic acid (Sigma, ref. 07260) and 50 mM Bis-Tris⋅HCl (pH 7.0). To solubilize mitochondrial membranes, the resuspended pellet was treated with digitonin (Sigma, ref. D141-500mg) at a final concentration of 6 mg per gram of starting sample and incubated at 4°C for 10 min. After centrifugation at 13,000 *× g* for 30 min, the supernatant containing solubilized mitochondrial proteins was carefully collected, and protein concentration was quantified using the BCA Protein Assay kit (Thermo Fisher Scientific, ref. 23227). For assessment of respiratory complex I assembly, digitonin-solubilized mitochondrial samples (10-15 μg of protein) were loaded onto NativePAGE 3–12% Bis-Tris polyacrylamide gels (Invitrogen, ref. BN2012BX10). Following electrophoretic separation, proteins were transferred and analyzed by Western blotting using an antibody directed against the NDUFS1 subunit.

### Quantification of mitochondrial ROS, mitochondrial iron and lipid peroxidation by flow cytometry

Mitochondrial ROS, mitochondrial iron, and lipid peroxidation levels were determined using the fluorescent probes MitoSox Red (Molecular Probes, ref. M36008), Mito-Ferro Green (Dojindo, ref. M489-10), and BODIPY™ 581/591 C11 (Thermo Fisher, ref. D3861), respectively. Following magnetic separation, isolated neurons and astrocytes (or non-neuronal cells) from the cerebrum, cerebellum, and DRG were incubated with 5 µM MitoSox Red, 10 µM MitoFerroGreen, or 10 µM BODIPY for 30 min at 37°C in DPBS supplemented with 0.5% BSA. After incubation, cells were centrifuged at 700 *× g* for 5 min, resuspended in DPBS with 0.5% BSA, and analyzed by flow cytometry. Samples stained with MitoSox Red or MitoFerroGreen were analyzed on a BD FACS Canto II flow cytometer (Becton Dickinson), whereas samples stained with BODIPY™ 581/591 C11 were analyzed on a CytoFlex SRT flow cytometer (Beckman Coulter). For all samples, instrument settings and the target cell populations were established using unstained controls, and debris was excluded from the analysis based on FSC/SSC light scattering profiles.

### Western blotting and antibodies

Cell extracts were prepared from isolated neurons and astrocytes or non-neuronal cells from the cerebrum, cerebellum, and DRG by sonication, followed by heating at 95 °C in lysis buffer containing 2% SDS, 125 mM Tris-HCl (pH 7.4), protease inhibitor cocktail (Roche, ref. 04693159001), and phosphatase inhibitor cocktail (Roche, ref. 04906845001). Protein concentration was determined using the BCA assay. A total of 10-15 µg of protein was subjected to SDS-polyacrylamide gel electrophoresis (SDS-PAGE) and transferred to PVDF membranes, except for frataxin, which was transferred onto nitrocellulose membranes. All membranes were blocked in 0.3% I-Block (Thermo Fisher Scientific, ref. T2015) and 0.1% Tween-20 (v/v) (Sigma-Aldrich, ref. P7949) in phosphate buffered saline (PBS). Protein extracts from digitonin-solubilized mitochondria were resolved in BNGE (as detailed in above), followed by transfer to PVDF membranes (300 mA at 60 V at 4°C). These membranes were blocked with 5% non-fat dry milk in TBST (20 mM Tris-HCl pH 8.0, 125 mM NaCl, 0.1% Tween-20 (v/v)). The primary antibodies employed were: Microtubule Associated Protein 2 (MAP2; 1:1000, Sigma-Aldrich, ref. M4403), Glial Fibrillary Acidic Protein (GFAP; 1:1000, Santa Cruz Biotechnology, ref. sc-33673), NADH:Ubiquinone Oxidoreductase Core Subunit S1 (NDUFS1; 1:1000, Abcam, ref. ab169540), ATP Synthase Peripheral Stalk Subunit B (ATPβ; 1:1000, Cell Signaling Technology, ref. 1672532S), Transferrin Receptor 1 (TFR1; 1:500, Invitrogen, ref. 13-6800), Ferritin Heavy Chain 1 (FTH1; 1:1000, Abcam, ref. ab75973), Glutathione Peroxidase 4 (GPX4; 1:1000, Santa Cruz Biotechnology, ref. sc-166570), NRF2 (1:1000, Abcam, ref. ab62352), and frataxin (1:1000, Abcam, ref. ab219414). Following immunoblotting, PVDF membranes were stained with Coomassie Brilliant Blue (CBB) and nitrocellulose membranes with Colloidal Gold (Bio-Rad, ref. 170-6527). Total protein staining was quantified using Image Lab software (Bio-Rad Laboratories) and used as the loading control for normalization of target protein levels. For digitonin-solubilized mitochondrial samples analyzed by BNGE, ATPβ was used as the loading control.

### Immunofluorescence, image acquisition and analysis

DRG tissue was processed following a previously established protocol (22). Briefly, mice were anesthetized via intraperitoneal injection of a ketamine/xylazine solution (100/10 mg/kg) followed by transcardial perfusion with 0.9% saline and subsequently with 4% paraformaldehyde in PBS. DRGs were then dissected and post-fixed in the same fixative overnight at 4°C. The dissected DRGs were cryoprotected by immersion in 30% sucrose in 100 mM phosphate buffer for 24 h at 4°C. Cryoblocks were prepared by embedding the cryoprotected ganglia in Tissue Freezing Medium (TFM, Electron Microscopy Sciences, ref. 72592) on dry ice. Cryosections were cut at a thickness of 14 μm using a freezing cryostat (Leica). For immunofluorescent staining, sections were permeabilized for 1 h in 0.3% Triton X-100 in PBS and subsequently blocked for 1 h in 5% BSA with 0.3% Triton X-100 in PBS. Sections were incubated overnight at 4°C with the following primary antibodies: GFAP (1:50, Santa Cruz Biotechnology, ref. sc-33673), Glutamine Synthetase (GS; 1:50, Abcam, ref. ab228590), TFR1 (1:50, Invitrogen, ref. 13-6800), FTH1 (1:50, Abcam, ref. ab75973), GPX4 (1:50, Santa Cruz Biotechnology, ref. sc-166570), NRF2 (1:50, Abcam, ref. ab62352), and frataxin (1:50, Abcam, ref. ab219414). After three washes with PBS, sections were incubated for 1 h at room temperature in the dark with the appropriate secondary antibodies: Alexa Fluor 488 goat anti-mouse (1:400, Fisher Scientific, ref. A11017) and Alexa Fluor 594 goat anti-rabbit (1:400, Invitrogen, ref. A21428). Sensory neurons were stained using NeuroTrace™ 435/455 Blue Fluorescent Nissl Stain (1:100, Life Technologies, ref. N21479). Finally, stained samples were washed and mounted with Mowiol (Calbiochem). Images were acquired using an Olympus FV1000 confocal microscope equipped with a 60× objective. Z-stack images were collected using identical laser intensity and gain settings across all experimental groups to ensure comparability. For each mouse (n = 3 per group), 1–2 DRG sections were analyzed, yielding a total population of approximately 500-2,000 cells per condition. Image quantification was performed using ImageJ software (National Institutes of Health, Bethesda, MD, USA). Neuronal regions were identified by Nissl staining, while SGCs were delineated using GFAP or GS immunostaining. Protein expression levels were quantified as fluorescence intensity per unit area, and particle analysis was used to compute mean intensity values for each region of interest.

### Statistical analysis

Data are presented as mean ± SEM. Normality was assessed via the Shapiro-Wilk test. Independent Student’s t-tests were used to evaluate the impact of the mutation (WT vs. I151F) within each specific cell type. Global comparisons across multiple groups or cell types were performed using one-way or two-way ANOVA, as appropriate, followed by Holm–Šídák or Tukey post hoc tests. For non-normal two-group datasets, the Mann–Whitney U test was used. Precise p-values are reported in each figure. No adjustments for multiple testing were applied. Representative images were selected to align with group averages. Statistical significance was defined as p < 0.05. All statistical analyses and graph generation were performed using GraphPad Prism 9.0 (GraphPad Software, Inc., La Jolla, CA).

## Results

### 1. Frataxin deficiency induces cell-type–specific alterations in mitochondrial respiration and supercomplexes organization in cerebrum neurons and astrocytes of FXN^I151F^ mice

Distinct cell populations from the cerebrum, cerebellum, and DRG of the FXN^I151F^ mouse model were purified using immunomagnetic separation. As detailed in the Materials and Methods, this approach enabled the isolation of neurons and astrocytes from both cerebrum and cerebellum. Neuronal populations were obtained by depleting non-neuronal cells using immunomagnetic antibodies targeting non-neuronal markers, whereas astrocytes were purified using immunomagnetic antibodies against ACSA-2 (Supplementary Fig. S1A). For DRG, since there is no specific isolation kit for SGCs is available, neurons were separated from the non-neuronal cell population using immunomagnetic antibodies directed against non-neuronal cells (Supplementary Fig. S1B). Following immunomagnetic separation, the purity of each cellular fraction from the cerebrum, cerebellum, and DRG was assessed by measuring the expression of MAP2, a neuronal marker, and GFAP, a glial marker (Fig. 1A). High MAP2 expression was detected in neuronal fractions from all three regions. In contrast, GFAP expression was markedly enriched in astrocyte fractions from the cerebrum and cerebellum. In the DRG, GFAP served as a glial marker in both Western blot and immunofluorescence analyses. Although, GFAP can label multiple glial subtypes, in the DRG it predominantly marks SGCs, the principal glial population. Accordingly, GFAP expression was restricted to the non-neuronal fraction in Western blot analysis (Fig. 1A), confirming effective neuronal–glial separation. Consistently, immunofluorescence in WT and FXN^I151F^ samples revealed GFAP and GS immunoreactivity surrounding Nissl-positive neuronal cell bodies, further validating SGC-specific localization (Fig. 1B).

**Figure 1.**
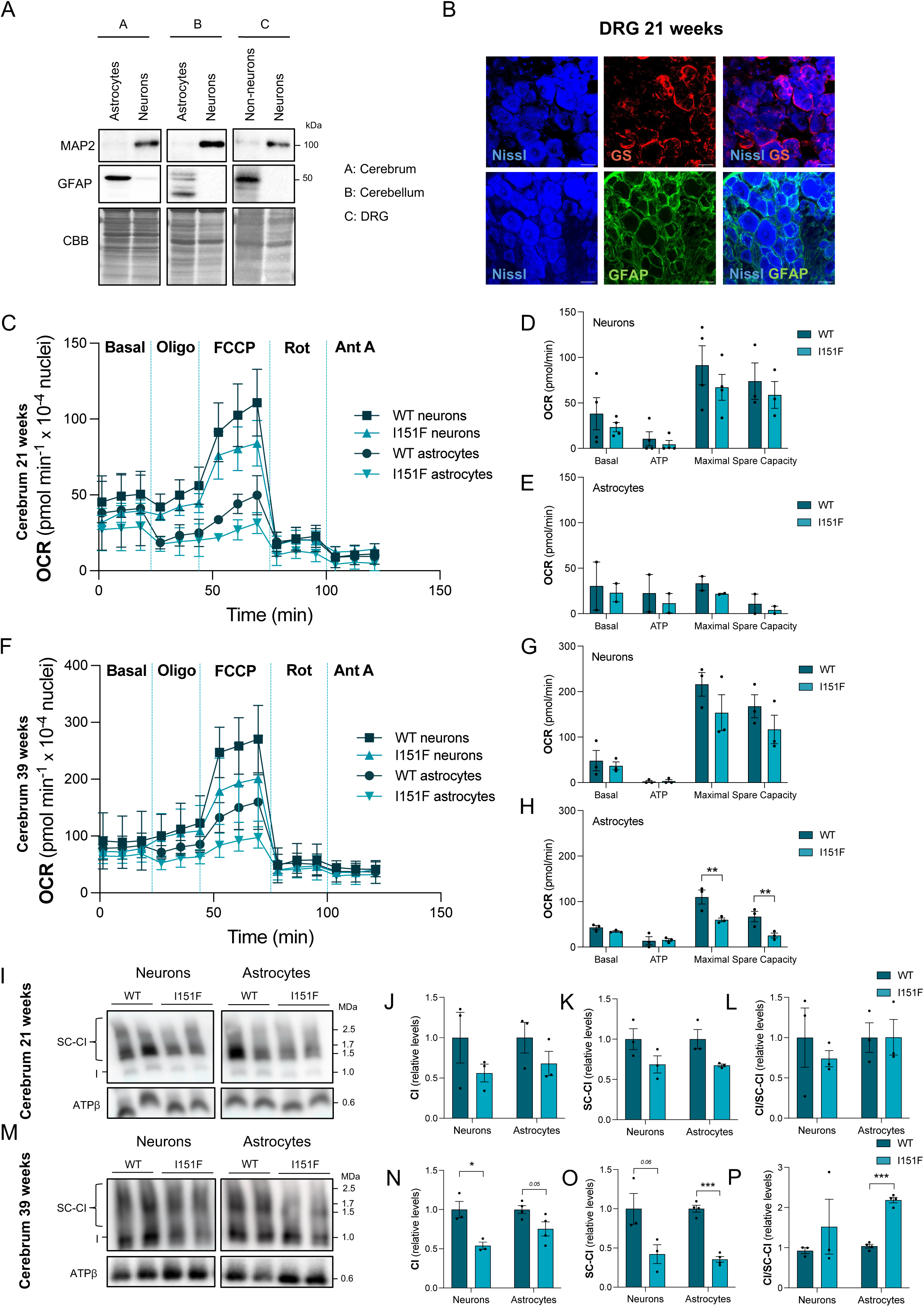
Cell-specific and age-dependent mitochondrial dysfunction in neurons and astrocytes from FXN^I151F^ mice model. A) Cell and Tissue Characterization: Western blot analysis of cell-type purity, showing expression of MAP2 (neuronal marker) and GFAP (glial marker) in distinct populations of astrocytes, neurons, and non-neuronal cells isolated from the cerebrum, cerebellum, and DRG of WT mice. CBB staining serves as a loading control. B) DRG Glial Markers: Immunohistochemical localization of glutamine synthetase (GS) and GFAP as satellite glial cell markers within DRG of 21-week-old mice. Representative images show Nissl staining (blue), GS (red), GFAP (green), and the composite image. Scale bar = 20 μm. C, F) Mitochondrial Respiration: Mitochondrial stress analysis of isolated cerebral neurons and astrocytes from WT and FXN^I151F^ mice at 21 weeks (C) and 39 weeks (F) of age, assessed by oxygen consumption rate (OCR) using the Seahorse XF Analyzer. D, E, G, H) Histograms representing the mean ± SEM for basal respiration, ATP-linked respiration, maximal respiration rate, and spare respiratory capacity in neurons (D, G; n=4 animals/group) and astrocytes (E, H; n=2-3 animals/group) at 21 weeks (D, E) and 39 weeks (G, H). I, M) Isolated mitochondria from cerebral neurons and astrocytes of WT and FXN^I151F^ mice at 21 weeks (I) and 39 weeks (M) were analyzed by Blue native-polyacrylamide gel electrophoresis (BN-PAGE) followed by immunoblotting. NDUFS1 (complex I) antibody was used for detection, with ATPβ serving as a loading control. J-L, N-P) Histograms representing the mean ± SEM (n=3 animals/group) for the relative quantification of free complex I (CI) content (J, N), total supercomplex (SC-CI) content (K, O), and the ratio of CI/SC-CI (L, P), derived from the NDUFS1 immunoblotting at 21 weeks (J-L) and 39 weeks (N-P). Significant differences between WT or FXN^I151F^ are indicated with p values <0.05(*), 0.01(**) or 0.001(***).

The molecular validity of the model was first confirmed by assessing frataxin protein levels in isolated cell populations. In WT mice, neurons and astrocytes from both the cerebrum and cerebellum exhibited robust and comparable frataxin expression across all tissues and ages examined (Supplementary Fig. S2A-H), although WT astrocytes in the cerebrum showed significantly higher frataxin levels than neurons at 39 weeks (Supplementary Fig. S2D). In contrast, FXN^I151F^ mice exhibited a severe and significant reduction in frataxin expression across all analyzed cell types and regions. In the cerebrum and cerebellum, frataxin levels were markedly decreased in both neurons and astrocytes at 21 and 39 weeks of age (Supplementary Fig. S2A-H). In the DRG, due to the low yield obtained after immunoseparation of cellular fractions, frataxin expression was assessed by immunofluorescence. Neurons were identified by Nissl staining, whereas SGCs were identified based on their characteristic anatomical localization, closely enveloping the neuronal somata and by their expression of GS and GFAP. In this tissue, frataxin levels were significantly reduced in both Nissl-positive neurons and GFAP-positive SGCs compared to WT mice (Supplementary Fig. S3A and B). These results confirm that the FXN^I151F^ mutation leads to a severe and uniform loss of frataxin across neuronal and glial populations throughout both the CNS and PNS.

To investigate how frataxin deficiency affects metabolic function in distinct neural cell types, mitochondrial bioenergetics were assessed in neuronal and astrocytic populations isolated from the cerebrum of WT and FXN^I151F^ mice at 21 and 39 weeks of age. Given the fundamental metabolic differences between neurons and astrocytes, particularly the reliance of neurons on OXPHOS for survival (72), OCR was quantified using Seahorse XF technology. Across both ages examined, OCR was consistently higher in neurons than in astrocytes (Fig. 1C-H). At 21 weeks of age, no differences in mitochondrial respiration were detected between FXN^I151F^ and WT mice in either neurons or astrocytes (Fig. 1D and E). By 39 weeks, a clearer pattern emerged. Neurons from FXN^I151F^ mice showed reductions in maximal respiration (∼29%) and spare respiratory capacity (∼31%), although these decreases did not reach statistical significance (Fig. 1G). In contrast, astrocytes from FXN^I151F^ mice displayed a significant decline in both maximal respiration (∼46%) and spare respiratory capacity (∼62%) relative to WT astrocytes (Fig. 1H).

Previous studies have reported cell-type–specific differences in the organization of mitochondrial supercomplexes, with astrocytes exhibiting a higher proportion of free complex I relative to complex I super-assembled into respiratory supercomplexes, a configuration associated with low bioenergetic efficiency and high ROS production (72,73). Based on these observations, the structural organization of the mitochondrial respiratory chain was examined in neurons and astrocytes isolated from the cerebrum of FXN^I151F^ mice (Fig. 1I-P). To evaluate mitochondrial respiratory chain super-assembly, BNGE was performed on digitonin-solubilized mitochondria from isolated neuronal and astrocytic populations. Native protein complexes were analyzed by immunoblotting using an antibody against NDUFS1, a complex I subunit detected that appears in both its monomeric (free) form and within I+III₂ and I₂+III₂ supercomplexes (75) (Fig. 1I and M). Consistent with previously published findings with whole tissue (20,21), FXN^I151F^ cells exhibited an overall reduction in complex I levels in both neurons and astrocytes. At 21 weeks of age, although levels of both free and supercomplex–associated complex I trended downward, the ratio of free–to–assembled complex I remained unchanged in both cell types (Fig. 1J-L). By 39 weeks, however, age–dependent alterations in supercomplexes organization became evident. In neurons, decreases in both free complex I and supercomplex–assembled complex I persisted, maintaining a relatively stable ratio (Fig. 1N-P). In contrast, 39–week–old FXN^I151F^ astrocytes displayed a more pronounced disruption of supercomplex stability: although both forms of complex I were reduced, the substantial loss of supercomplexes (Fig. 1O) resulted in a significant increase in the free–to–assembled complex I ratio (Fig. 1P). These findings suggest that while complex I depletion is a general feature, the failure to incorporate complex I into supercomplexes is specifically exacerbated in astrocytes at both ages analysed.

### 2. Increased mitochondrial ROS and iron overload in FXN^I151F^ astrocytes and DRG cells

The observed reduction in supercomplex assembly in FXN^I151F^ mice, together with the established role of complex I as a major source of mitochondrial ROS (73), might contribute to increased oxidative stress in the mutant. To test this, mitochondrial ROS levels were quantified in neuronal and astrocytic populations from the cerebrum and cerebellum, as well as in neuronal and non-neuronal cells from the DRG, at 21 and 39 weeks of age. Quantification was performed by flow cytometry using the MitoSox Red probe following immunomagnetic separation of each cell population (Fig. 2A-F; Supplementary Fig. S4). In both the cerebrum and cerebellum, MitoSox fluorescence, reflecting mitochondrial ROS levels, was consistently higher in WT astrocytes than in WT neurons across both ages (Fig. 2A, B, D and E). At 21 weeks, neuronal ROS levels did not differ between FXN^I151F^ and WT mice in either region, whereas FXN^I151F^ astrocytes exhibited increased mitochondrial ROS relative to WT astrocytes (Fig. 2A and B). In the DRG, both neurons and non-neuronal cells from FXN^I151F^ mice exhibited elevated mitochondrial ROS compared to with WT cells (Fig. 2C). At 39 weeks, neuronal ROS levels in the cerebrum remained unchanged, while astrocytes from FXN^I151F^ mice showed a non-significant upward trend relative to WT astrocytes (Fig. 2D). In contrast, both neurons and astrocytes in the cerebellum, as well as neurons and non–neuronal cells in the DRG, exhibited clear increases in mitochondrial ROS in FXN^I151F^ mice relative to WT (Fig. 2E and F). The elevations in mitochondrial ROS observed in specific cell populations of the FXN^I151F^ mouse model, such as cerebellar astrocytes and both neuronal and non-neuronal DRG cells, strongly indicate increased mitochondrial oxidative stress within these compartments. Redox–active metal ions, particularly iron, are major contributors to mitochondrial ROS production (81). Previous evidence has established a link between frataxin deficiency and disrupted iron homeostasis (87–92), a finding corroborated in the FXN^I151F^ mouse model (22,85). Although the neurotoxic potential of iron has been studied, many prior studies have not distinguished iron dysregulation across specific neural cell types, potentially overlooking differences in iron metabolism between neurons, astrocytes, and other glial cells. To address this, mitochondrial Fe²⁺ levels were quantified in purified neuronal and glial populations from the cerebrum, cerebellum, and DRG using the Mito–FerroGreen probe and flow cytometry after immunomagnetic separation (Fig. 2G–L; Supplementary Fig. S5). In both the cerebrum and cerebellum, WT astrocytes consistently exhibited higher mitochondrial Fe²⁺ content than WT neurons at both ages examined (Fig. 2G, H, J, K). At 21 weeks, mitochondrial Fe²⁺ levels in cerebrum did not differ between FXN^I151F^ and WT mice, neither in neurons or astrocytes (Fig. 2G). In the cerebellum, neuronal mitochondrial Fe²⁺ levels remained unchanged, whereas cerebellar astrocytes exhibited increased mitochondrial Fe²⁺ in FXN^I151F^ mice compared with WT (Fig. 2H). In the DRG, both neurons and non-neuronal cells from FXN^I151F^ mice displayed significantly elevated mitochondrial Fe²⁺ relative to WT (Fig. 2I). At 39 weeks, neuronal mitochondrial Fe²⁺ levels remained unchanged between genotypes in both cerebrum and cerebellum, whereas astrocytes in these regions showed increased mitochondrial Fe²⁺ relative to WT astrocytes (Fig. 2J and K). Similarly, in the DRG, both neurons and non-neuronal cells from FXN^I151F^ mice exhibited higher mitochondrial Fe²⁺ levels than their respective WT cells at this age (Fig. 2L).

**Figure 2.**
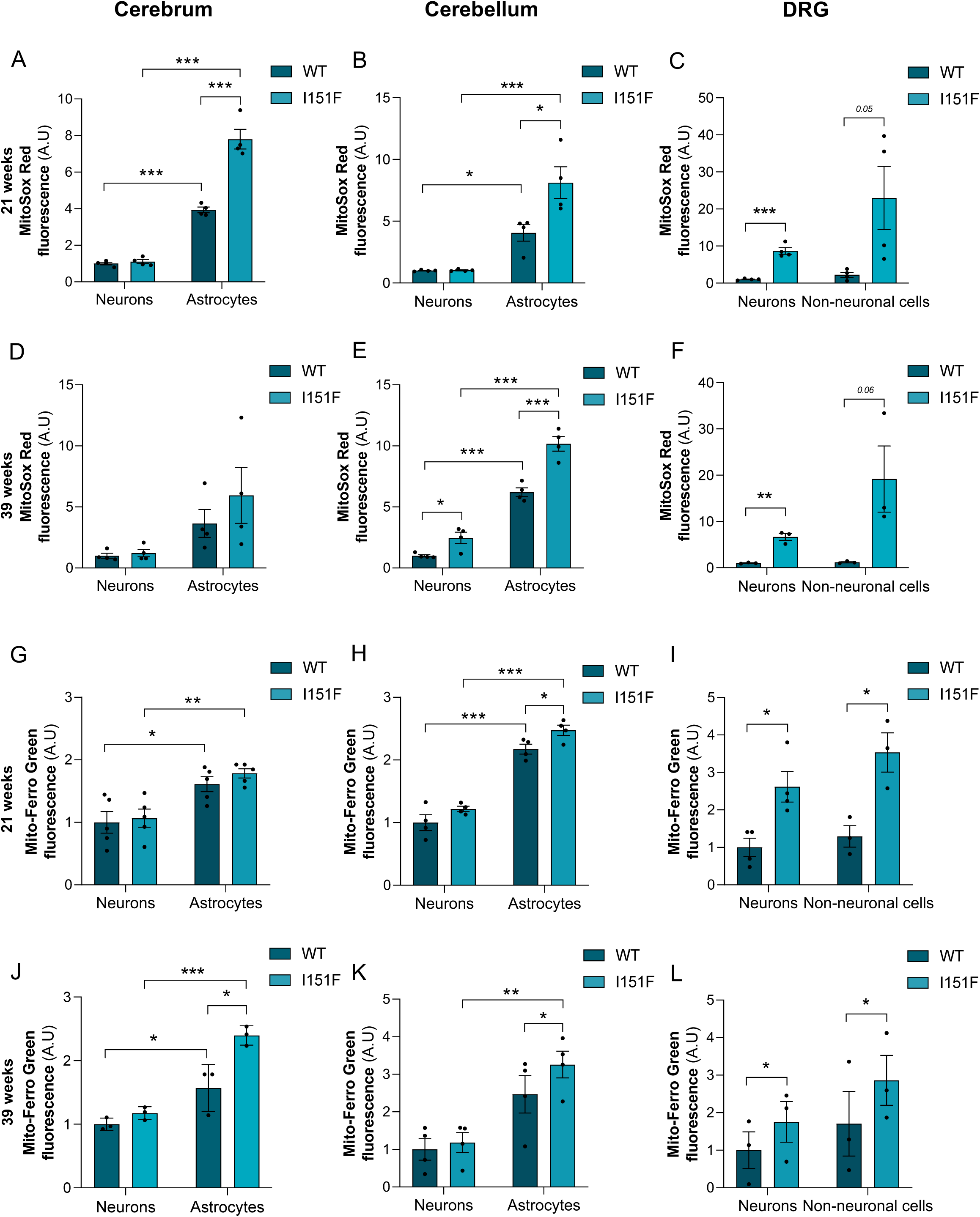
Mitochondrial ROS and mitochondrial iron (Fe^2+^) in distinct neuronal and glial populations of FXN^I151F^ mice. Mitochondrial ROS and Fe^2+^ were quantified using flow cytometry with the MitoSox Red and Mito-Ferro Green probes, respectively. Cell populations analyzed include neurons and astrocytes (cerebrum, cerebellum) or neurons and non-neuronal cells (DRG). A-F) Mitochondrial ROS (MitoSox Red): Histograms represent the mean ± SEM of MitoSox Red fluorescence quantification at 21 weeks of age (Cerebrum: A, Cerebellum: B, DRG: C; n=4 animals/group) and 39 weeks of age (Cerebrum: D, Cerebellum: E, DRG: F; n=3-4 animals/group). G-L) Mitochondrial Fe^2+^ (Mito-Ferro Green): Histograms represent the mean ± SEM of Mito-Ferro Green fluorescence quantification at 21 weeks of age (Cerebrum: G, n=5; Cerebellum: H, n=4; DRG: I, n=3 animals/group) and 39 weeks of age (Cerebrum: J, n=3; Cerebellum: K, n=4; DRG: L, n=3 animals/group). Significant differences between WT or FXN^I151F^ are indicated with p values <0.05(*), 0.01(**) or 0.001(***).

### 3. Cell-type specific iron dyshomeostasis in the FXN^I151F^ mouse model

Frataxin deficiency is known to promote mitochondrial iron accumulation and disrupt systemic iron homeostasis. Previous studies have reported increased TFR1 and decreased FTH1 in the DRG (22), as well as elevated FTH1 in the cerebellum of 21-week-old FXN^I151F^ mice (85). However, whether these alterations differ across specific neural cell types—particularly neurons versus astrocytes in the cerebrum and cerebellum—remains unclear. As shown in Fig. 2G-L, astrocytes generally exhibit higher mitochondrial Fe²⁺ levels than neurons under physiological conditions, and this difference is exacerbated by frataxin-deficiency. To investigate cell–type–specific iron handling, TFR1 (iron uptake) and FTH1 (iron storage) protein levels were quantified by Western blot in isolated neuronal and astrocytic populations from the cerebrum and cerebellum of FXN^I151F^ mice. In the cerebrum, TFR1 levels in both neurons and astrocytes showed no significant differences at 21 weeks of age (Fig. 3A and B). By 39 weeks, TFR1 levels remained unchanged in cerebral neurons, whereas FXN^I151F^ astrocytes exhibited a significant decrease compared with WT astrocytes (Fig. 3C and D). In the cerebellum, a more complex pattern was observed. At 21 weeks, FXN^I151F^ neurons displayed decreased TFR1 levels relative to WT neurons, while FXN^I151F^ astrocytes showed increased TFR1 levels compared to WT astrocytes (Fig. 3E and F). By 39 weeks, cerebellar neurons from mutant mice maintained TFR1 levels comparable to WT, whereas FXN^I151F^ astrocytes exhibited markedly higher TFR1 levels relative to WT (Fig. 3G and H). FTH1 levels were assessed in parallel (Fig. 3I-P). In the cerebrum at 21 weeks, both neurons and astrocytes from FXN^I151F^ mice showed increased FTH1 levels compared to WT (Fig. 3I and J). At 39 weeks, neuronal FTH1 levels remained unchanged, while astrocytic FTH1 was significantly reduced compared with WT astrocytes (Fig. 3K and L). In the cerebellum, at 21 weeks, FXN^I151F^ neurons exhibited increased FTH1 levels, whereas FXN^I151F^ astrocytes showed decreased FTH1 relative to WT cells (Fig. 3M and N). By 39 weeks, neuronal FTH1 levels remained elevated, and astrocytic FTH1 levels decreased further compared with WT astrocytes (Fig. 3O and P). TFR1 and FTH1 levels were assessed in DRG from FXN^I151F^ mice at 21 weeks using immunofluorescence staining in both neuronal and non-neuronal cells. Nissl-positive neurons exhibited a 6-fold increase in TFR1 levels relative to WT neurons, whereas GFAP-positive SGCs showed a 2.2-fold increase relative to WT SGCs (Fig. 4A and B). In contrast, FTH1 levels were markedly reduced, with a 5-fold decrease in neurons and a 7-fold decrease in SGCs (Fig. 4C and D).

**Figure 3.**
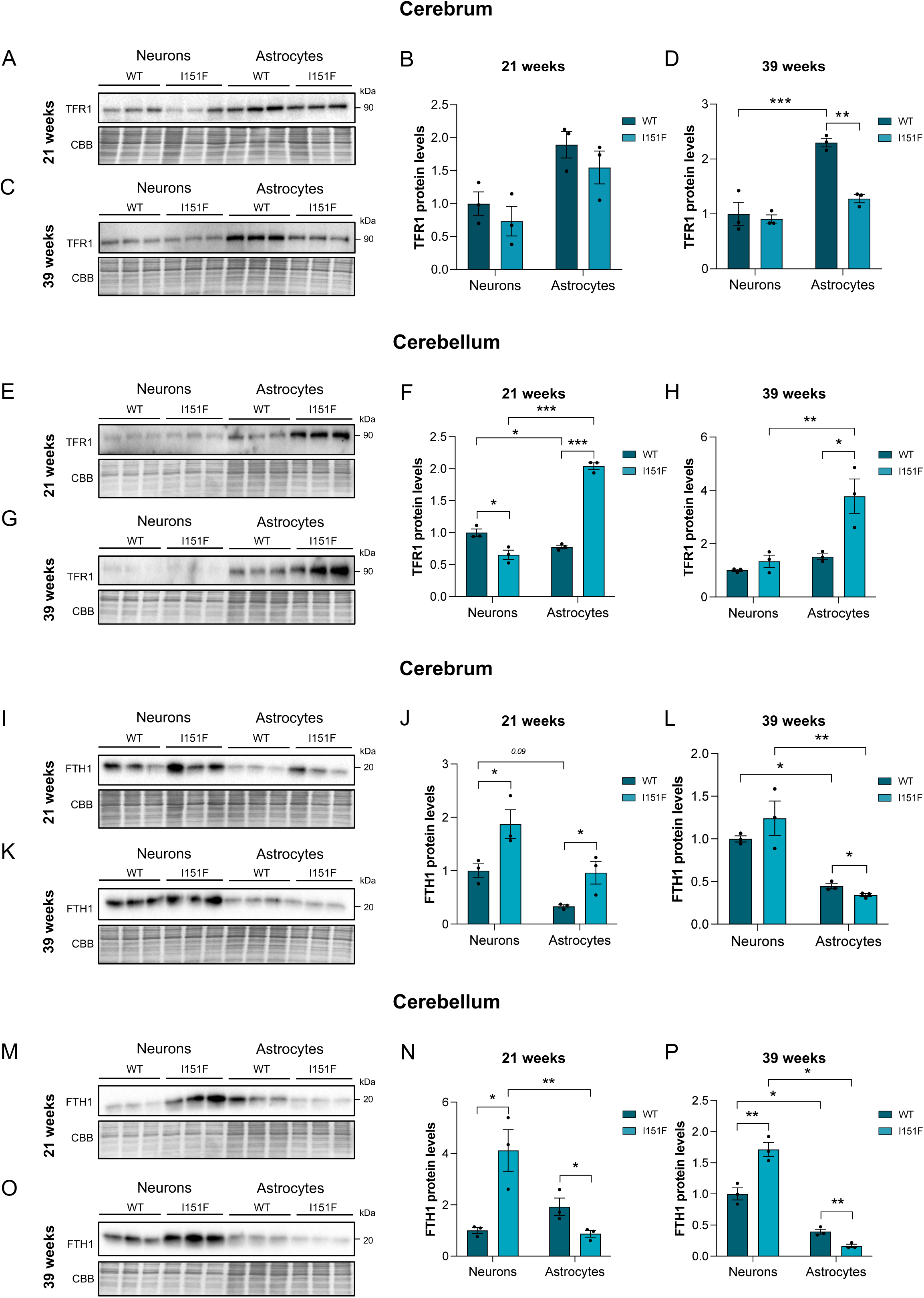
Iron uptake (TFR1) and storage (FTH1) levels in cell-specific populations from FXN^I151F^ mice. Protein levels of the iron uptake regulator Transferrin Receptor 1 (TFR1) and the iron storage protein Ferritin Heavy Chain 1 (FTH1) were analyzed by Western blotting in isolated neurons and astrocytes from the cerebrum and cerebellum of control WT and FXN^I151F^ mice. CBB staining was used as a loading control for all blots. A, B, E, F) Representative Western blots showing TFR1 protein levels in neurons and astrocytes isolated from the cerebrum (A: 21 weeks; B: 39 weeks) and cerebellum (E: 21 weeks; F: 39 weeks). C, D, G, H) Corresponding histograms representing the mean ± SEM (n=3 mice per group) of TFR1 protein quantification for the cerebrum (C: 21 weeks; D: 39 weeks) and cerebellum (G: 21 weeks; H: 39 weeks). I, J, M, N) Representative Western blots showing FTH1 protein levels in neurons and astrocytes isolated from the cerebrum (I: 21 weeks; J: 39 weeks) and cerebellum (M: 21 weeks; N: 39 weeks). K, L, O, P) Corresponding histograms representing the mean ± SEM (n=3 mice per group) of FTH1 protein quantification for the cerebrum (K: 21 weeks; L: 39 weeks) and cerebellum (O: 21 weeks; P: 39 weeks). Significant differences between WT or FXN^I151F^ are indicated with p values <0.05(*), 0.01(**) or 0.001(***).

**Figure 4.**
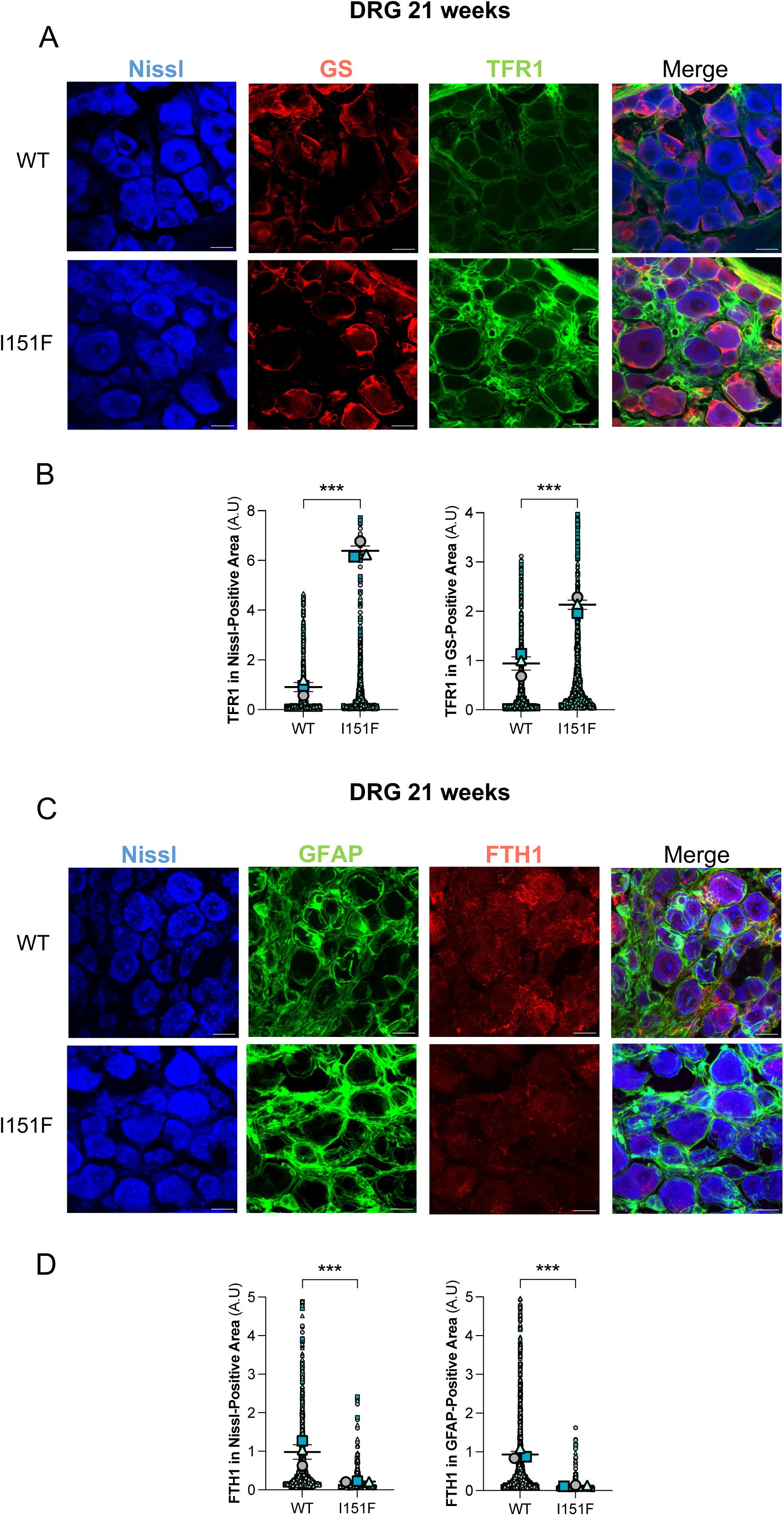
Iron uptake and storage in neurons and non-neuronal cells isolated from DRG FXN^I151F^ mice. Iron uptake (TFR1) and storage (FTH1) protein levels were assessed via immunofluorescence in DRG sections. Nissl staining (blue) was used to identify neurons. SGCs (non-neuronal cells) were identified using Glutamine Synthetase (GS, red) or Glial Fibrillary Acidic Protein (GFAP, green). The scale bar for all images is 20 μm. A) Representative immunofluorescence images showing TFR1 in green and its localization with the cell markers. B) Quantification of TFR1 immunofluorescence intensity in Nissl-positive neurons and GS-positive satellite cells. C) Representative immunofluorescence images showing FTH1 in red and its localization with the cell markers. D) Quantification of FTH1 immunofluorescence intensity in Nissl-positive neurons and GFAP-positive satellite cells. All quantification data (B and D) are presented as the mean ± SEM (n=3 independent mice). Each biological replicate is color-coded, with individual data points representing different cells/particles analyzed per experiment. Significant differences between WT or FXN^I151F^ are indicated with p values <0.05(*), 0.01(**) or 0.001(***).

### 4. Ferroptosis markers in glial cells of the FXN^I151F^ mouse model

Given the established relationship between ferroptosis and frataxin deficiency (22,93), we next investigated ferroptotic cell death, a lipid-peroxidation-driven process triggered by the oxidation of plasma membrane polyunsaturated fatty acids (PUFAs). Excess free iron catalyses the generation of highly reactive radicals via the Fenton reaction, leading to oxidative stress and lipid peroxidation. To evaluate ferroptosis in the FXN^I151F^ mouse model, lipid peroxidation was quantified in neuronal and glial cell populations from the cerebrum, cerebellum, and DRG using flow cytometry with the BODIPY C11 581/591 probe following immunomagnetic cell separation. In the cerebrum at 21 weeks, the oxidized/reduced BODIPY-C11 ratio did not differ significantly between FXN^I151F^ and WT mice in either neurons or astrocytes (Fig. 5A). By 39 weeks, neuronal lipid peroxidation levels remained unchanged between FXN^I151F^ and WT mice, whereas FXN^I151F^ astrocytes exhibited a marked increase compared to WT astrocytes (Fig. 5B). Similarly, in the cerebellum, no differences were observed at 21 weeks in either neurons or astrocytes (Fig. 5C); however, at 39 weeks, lipid peroxidation was significantly elevated in FXN^I151F^ astrocytes relative to WT astrocytes, while neuronal levels remained unaffected (Fig. 5D). In the DRG at 21 weeks, both neurons and non-neuronal cells from FXN^I151F^ mice displayed increased oxidized/reduced BODIPY-C11 ratios, with the elevation being more pronounced in non-neuronal cells (Fig. 5E). At 39 weeks, the increase remained significant only in non-neuronal cells, relative to WT (Fig. 5F).

**Figure 5.**
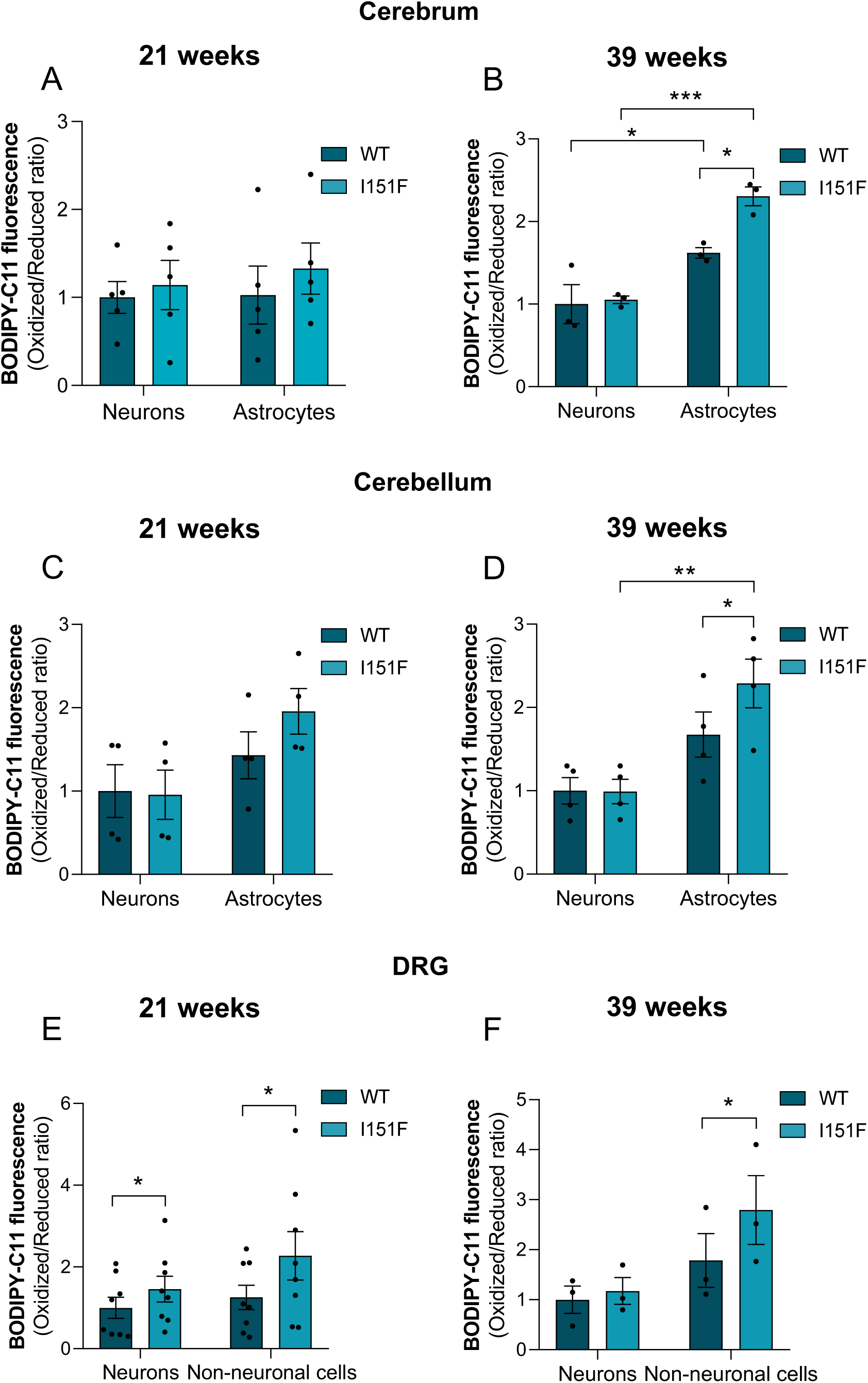
Lipid peroxidation in cell-specific populations isolated from the cerebrum, cerebellum, and DRG of FXN^I151F^ mice. Lipid peroxidation was assessed by flow cytometry using the BODIPY C11 probe. A), B), C), D), E) and F) The histograms represent the mean ± SEM of the ratio of oxidized (510 nm) to reduced (590 nm) BODIPY-C11 fluorescence in neurons and astrocytes isolated from the cerebrum (A-B) and cerebellum (C-D), and in neurons and non-neuronal cells from the DRG (E-F) of control WT and FXN^I151F^ mice at 21 weeks (A, C, and E) and 39 weeks (B, D, and F) of age. Significant differences between WT or I151F are indicated with p values <0.05(*), 0.01(**) or 0.001(***).

To further elucidate the molecular mechanisms underlying the increased lipid peroxidation observed across distinct cell populations in FXN^I151F^ mice, the expression of GPX4 was assessed in affected tissues and cell types. GPX4 is a key regulator of ferroptosis that detoxifies lipid hydroperoxides and protects membranes from oxidative damage. Overall, GPX4 levels were lower in astrocytes than in neurons (Fig. 6). In the cerebrum, GPX4 levels in FXN^I151F^ neurons did not differ from WT neurons at either 21 or 39 weeks of age (Fig. 6A-D). In contrast, FXN^I151F^ astrocytes exhibited significantly reduced GPX4 levels relative to WT astrocytes at both time points (Fig. 6A-D). In the cerebellum, FXN^I151F^ neurons displayed increased GPX4 levels at 21 weeks compared to WT neurons (Fig. 6E and F); however, by 39 weeks, neuronal GPX4 levels no longer differed between genotypes (Fig. 6G and H). Notably, cerebellar astrocytes from FXN^I151F^ mice demonstrated a consistent reduction in GPX4 expression at both 21 and 39 weeks relative to WT astrocytes (Fig. 6E-H). GPX4 levels were also analyzed in the DRG at 21 weeks of age using immunofluorescence staining. In FXN^I151F^ DRG, Nissl-positive neurons exhibited an approximately 5-fold decrease in GPX4 levels compared to WT neurons, while GFAP-positive SGCs showed an approximately 5.5-fold decrease relative to WT SGCs (Fig. 6I and J). These data indicate that GPX4 downregulation is a prominent feature of glial populations—particularly astrocytes and SGCs—under frataxin-deficient conditions and likely contributes to the cell-type–specific susceptibility to lipid peroxidation and ferroptotic stress.

**Figure 6.**
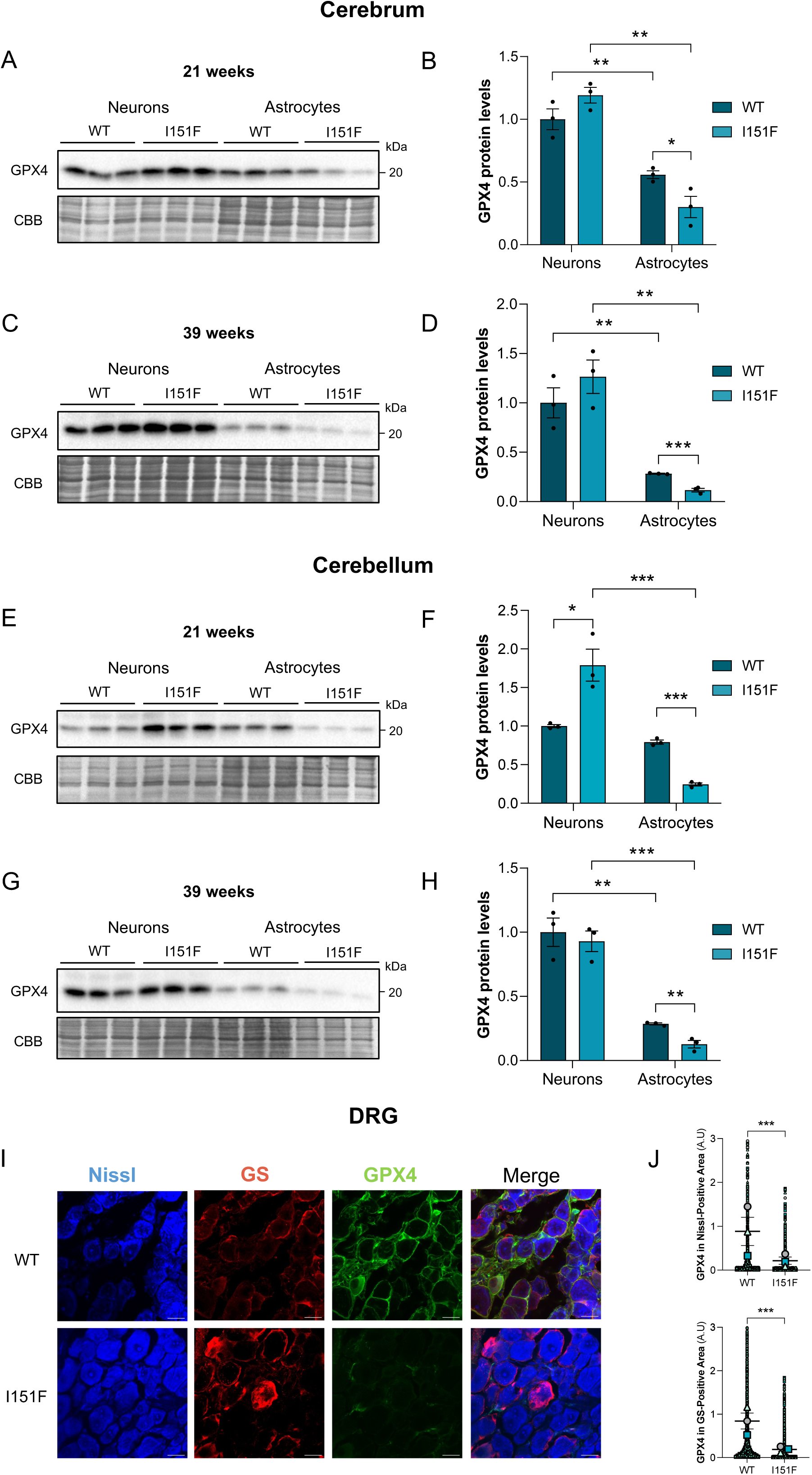
GPX4 Levels in cell-specific populations of FXN^I151F^ mice. Glutathione Peroxidase 4 (GPX4) protein levels were assessed in isolated neurons and astrocytes from the cerebrum and cerebellum by Western blotting, and in neurons and non-neuronal cells (satellite cells) from the DRG by immunofluorescence. A, C, E, G) Representative Western blots showing GPX4 protein levels in neurons and astrocytes isolated from the cerebrum (A: 21 weeks; C: 39 weeks) and cerebellum (E: 21 weeks; G: 39 weeks) of control WT and FXN^I151F^ mice. CBB staining served as the loading control. B, D, F, H) Corresponding histograms representing the mean ± SEM (n=3 mice per group) of GPX4 protein quantification for the cerebrum (B: 21 weeks; D: 39 weeks) and cerebellum (F: 21 weeks; H: 39 weeks). I) Representative immunofluorescence images showing GPX4 (green) localization within DRG sections. Nissl staining (blue) identifies neurons, and GS (red) identifies satellite glial cells. Scale bar = 20 μm. J) Quantification of GPX4 immunofluorescence intensity in Nissl-positive neurons and GS-positive satellite cells. Data are presented as the mean ± SEM (n=3 independent mice). Each biological replicate is color-coded, and individual data points represent different cells/particles analyzed per experiment. Significant differences between WT or FXN^I151F^ are indicated with p values <0.05(*), 0.01(**) or 0.001(***).

### 5. Altered NRF2 expression across cell types and nervous system regions in the FXN^I151F^ mouse model

Given these observations, the role of NRF2—a master regulator of cytoprotective gene expression coordinating cellular defences against oxidative stress and ferroptosis—was next examined. Previous work demonstrated that DRG from FXN^I151F^ mice exhibit significantly reduced total NRF2 levels together with diminished nuclear localization, indicative of impaired NRF2 activation and transcriptional activity (22). Notably, astrocytes have been reported to express higher basal NRF2 levels than neurons, a feature thought to confer increased resistance to redox imbalance (94). To determine whether NRF2 expression is differentially affected across brain regions and specific cell populations in the FXN^I151F^ mouse model, NRF2 protein levels were quantified in neurons and astrocytes isolated from the cerebrum, cerebellum, and DRG at distinct stages of disease progression. Overall, across cerebrum and cerebellum, NRF2 levels were higher in astrocytes than in neurons (Fig. 7A-H). In the cerebrum, neurons from FXN^I151F^ mice exhibited an increase in NRF2 levels at 21 weeks of age compared with WT neurons (Fig. 7A and B), but no significant differences were observed by 39 weeks (Fig. 7C and D). In contrast, FXN^I151F^ astrocytes showed a trend toward decreased NRF2 expression at 21 weeks (Fig. 7A and B), which became significantly reduced at 39 weeks relative to WT astrocytes (Fig. 7C and D). In the cerebellum, FXN^I151F^ neurons presented a consistent decline in NRF2 levels at both 21 and 39 weeks (Fig. 7E-H). FXN^I151F^ astrocytes from the cerebellum did not display statistically significant changes in NRF2 expression at 21 weeks (Fig. 7E and F), but showed a significant decrease by 39 weeks (Fig. 7G and H). NRF2 expression was also assessed in the DRG at 21 weeks of age using immunofluorescence staining. In FXN^I151F^ DRG, Nissl-positive neurons exhibited an approximately 2.9-fold reduction in NRF2 levels compared with WT neurons, while GFAP-positive SGCs showed an approximately 2.3-fold decrease relative to WT SGCs (Fig. 7I and J).

**Figure 7.**
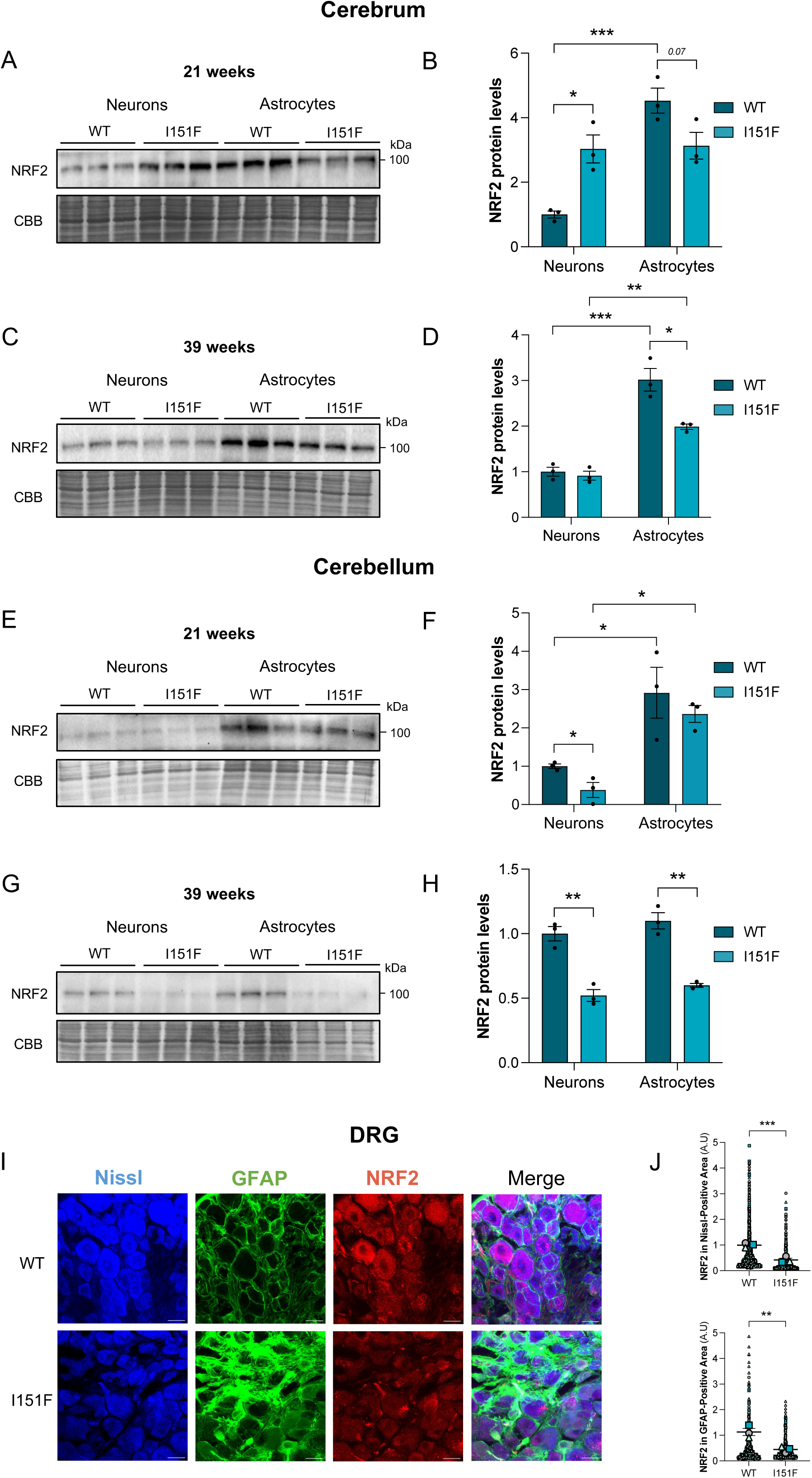
NRF2 levels in cell-specific populations of FXN^I151F^ mice. Nuclear factor erythroid 2-related factor 2 (NRF2) protein levels were assessed in isolated neurons and astrocytes from the cerebrum and cerebellum by Western blotting, and in neurons and non-neuronal cells (satellite cells) from the DRG by immunofluorescence. A, C, E, G) Representative Western blots showing NRF2 protein levels in neurons and astrocytes isolated from the cerebrum (A: 21 weeks; C: 39 weeks) and cerebellum (E: 21 weeks; G: 39 weeks) of control WT and FXN^I151F^ mice. CBB staining served as the loading control. B, D, F, H) Corresponding histograms representing the mean ± SEM (n=3 mice per group) of NRF2 protein quantification for the cerebrum (B: 21 weeks; D: 39 weeks) and cerebellum (F: 21 weeks; H: 39 weeks). I) Representative immunofluorescence images showing NRF2 (red) localization within DRG sections. Nissl staining (blue) identifies neurons, and GFAP (green) identifies satellite glial cells. Scale bar = 20 μm. J) Quantification of NRF2 immunofluorescence intensity in Nissl-positive neurons and GFAP-positive satellite cells. Data are presented as the mean ± SEM (n=3 independent mice). Each biological replicate is color-coded, and individual data points represent different cells/particles analyzed per experiment. Significant differences between WT or FXN^I151F^ are indicated with p values <0.05(*), 0.01(**) or 0.001(***).

### 6. Astrogliosis and satellite glial cell activation in the FXN^I151F^ mouse model

Astrocytes respond to neuronal damage and the associated release of toxic mediators by entering a reactive state—astrogliosis—typically marked by robust upregulation of GFAP (95). Since NRF2 plays a central role in regulating antioxidant defences and cytoprotective responses, the reduction in NRF2 levels observed in FXN^I151F^ astrocytes may compromise their ability to counteract oxidative stress, thereby exacerbating astrocyte activation and GFAP upregulation. To investigate this possibility, GFAP expression was analyzed in astrocytes isolated from the cerebrum and cerebellum, as well as in glial populations of the DRG. Similar to CNS astrocytes, SGCs undergo a reactive transformation in response to neuronal injury or pathological stress, a process termed “satellite gliosis” (40,46). The results revealed a significant increase in GFAP levels in astrocytes from both the cerebrum and cerebellum of FXN^I151F^ mice compared with WT controls at both 21 and 39 weeks of age (Fig. 8A-H). The most pronounced elevation occurred at 21 weeks (Fig. 8B and F), suggesting an early and robust gliotic response. In the DRG, immunofluorescence analysis revealed that GFAP expression was significantly upregulated in SGCs surrounding neuronal somata, with a 2.5-fold increase in FXN^I151F^ mice compared to WT at 21 weeks of age (Fig. 8I and J).

**Figure 8.**
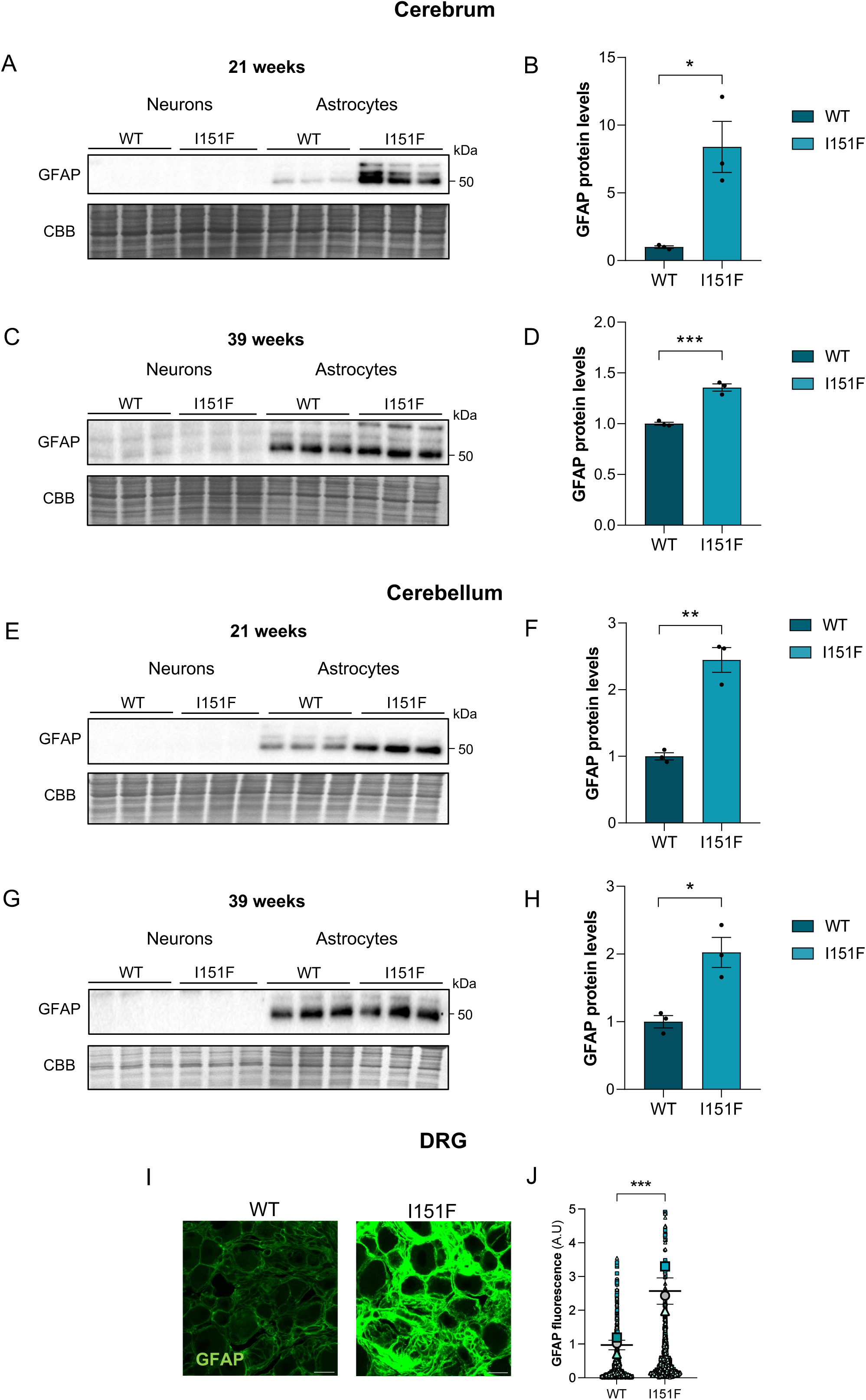
Astrogliosis (GFAP Levels) in the cerebrum, cerebellum, and DRG of FXN^I151F^ mice. Glial Fibrillary Acidic Protein (GFAP) levels, a marker of astrogliosis, were assessed in isolated astrocytes from the cerebrum and cerebellum by Western blotting, and in DRG sections by immunofluorescence. A, C, E, G) Representative Western blots showing GFAP protein levels in astrocytes isolated from the cerebrum (A: 21 weeks; C: 39 weeks) and cerebellum (E: 21 weeks; G: 39 weeks) of control WT and FXN^I151F^ mice. CBB staining served as the loading control. B, D, F, H) Corresponding histograms representing the mean ± SEM (n=3 mice per group) of GFAP protein quantification for the cerebrum (B: 21 weeks; D: 39 weeks) and cerebellum (F: 21 weeks; H: 39 weeks). I) Representative immunofluorescence images showing GFAP (green) within DRG sections. Scale bar = 20 μm. J) Quantification of GFAP immunofluorescence intensity. Data are presented as the mean ± SEM (n=3 independent mice). Each biological replicate is color-coded, with individual data points representing analyzed cells. Significant differences between WT or FXN^I151F^ are indicated with p values <0.05(*), 0.01(**) or 0.001(***).

## Discussion

The present study addresses a critical gap in FA research: the reliance on bulk tissue analyses, which has likely masked cell-type-specific contributions to disease pathology. While extensive prior work provided a comprehensive overview of frataxin deficiency across affected systems—including the cerebellum, DRG, heart, pancreas, and liver (20–22,85,90,96–98)—these studies relied primarily on whole-tissue homogenates. As a result, the cellular and molecular mechanisms underlying systemic pathology remain incompletely understood. Although cell-based models like patient-derived fibroblasts or primary cultures have contributed valuable insights, these *in vitro* systems fail to fully recapitulate the complex cellular interactions and microenvironment present *in vivo* (21,22,80,93,99). Therefore, detailed *in vivo* assessments of cell-type-specific mitochondrial and bioenergetic dysfunction in FA have remained elusive.

Recent advances in high-throughput molecular tools, included single-cell RNA-sequencing, have begun to address key limitations of bulk-tissue approaches in FA research (100–102). Nevertheless, when applied to heterogeneous tissues, these methods inevitably average distinct metabolic and functional attributes of different cell types, such as oxidative neurons versus glycolytic glia (64,68,103). Furthermore, transcriptomic analyses provides only static snapshots of cellular states; capturing key hallmarks features of mitochondrial dysfunction—such as OCR, iron handling, and superoxide generation—requires dynamic live-cell functional assays. To overcome these constraints, the present study employed advanced immunomagnetic separation strategies to isolate highly purified populations of neurons, astrocytes, and non-neuronal cells from the cerebrum, cerebellum, and the DRG. Cell-type purity was confirmed by canonical markers, like MAP2 for neurons and GFAP for glial cells. Notably, this work represents the first successful isolation of DRG cell populations for such analyses in FA. Although the lack of specific antibodies precluded the direct isolation of SGCs, the high-purity separation of DRG neurons from non-neuronal populations—including Schwann cells and SGCs—enabled robust comparative analyses (104). The central finding of this study is that, despite a uniform reduction of frataxin levels across cell types, pathological outcomes are strikingly cell-type-specific. Early and severe mitochondrial dysfunction, iron dyshomeostasis, and collapse of antioxidant defenses are concentrated predominantly within glial populations—most prominently cerebellar astrocytes—as well as within both neurons and glial cells of the DRG. These findings reposition glial cells as active contributors to FA pathogenesis, driving disease progression by fostering a toxic microenvironment (23–27,30,80,105,106). Elucidating these cell-type-specific consequences is essential to understanding the selective vulnerability of sensory neurons, the neuropathological hallmark of FA (28).

To dissect these mechanisms, we compared the mitochondrial functions in isolated neurons and glial cells. As expected, neurons showed significantly higher OCR than astrocytes, reflecting their greater reliance on OXPHOS (61,64,72,107). However, bioenergetic failure was most pronounced in glia. By 39 weeks of age, FXN^I151F^ astrocytes displayed marked reductions in maximal respiration and spare respiratory capacity, whereas mitochondrial deficits in neurons remained comparatively modest. This glia-centered vulnerability, observed in the cerebrum, suggests that cumulative metabolic stress drives an age-dependent collapse of astrocytic bioenergetics, in agreement with previous reports of respiratory impairment in frataxin-deficient cerebellar astrocytes (29). Structural analyses using BNGE combined with immunoblotting for the NDUFS1 subunit provided a basis for these functional defects. While earlier studies reported reduced complex I subunits in bulk tissues (20,21), our study found that, in the cerebrum from the FNX^I151F^ mice, the ratio of free-to-assembled complex I increased significantly only in astrocytes at 39 weeks, indicating disrupted supercomplex stability (75). This is supported by evidence that human frataxin directly interacts with respiratory complexes I, II, and III to preserve their structural integrity (108). Because supercomplex organization is vital for electron transport efficiency and for limiting ROS production (74), its disorganization explains the heightened oxidative stress observed in astrocytes. In contrast, cerebral neurons from mutant mice exhibited only mild alterations in NDUFS1 distribution, suggesting greater metabolic resilience.

The mechanistic link between mitochondrial structural defects and cellular toxicity lies in an iron-driven oxidative stress cascade (90,109). Using MitoSox Red and Mito-FerroGreen, this study confirmed that WT astrocytes naturally possess higher basal levels of ROS and mitochondrial ferrous ions than neurons, reflecting their lower energetic efficiency (72,73,110). Frataxin deficiency exacerbates this intrinsic vulnerability. While previous studies suggested that cerebellar astrocytes are particularly sensitive to frataxin loss (29,84,111), the present work provides the first direct comparison between neurons and glia across both the CNS and PNS. At an early, pre-symptomatic stage (21 weeks), mitochondrial ROS levels were already significantly elevated in FXN^I151F^ cerebrum and cerebellar astrocytes, as well as across all DRG populations. By 39 weeks, oxidative stress expanded to cerebellar neurons; nonetheless, in the cerebrum, it remained strictly confined to astrocytes. Mitochondrial iron accumulation followed a similar temporal and spatial pattern, appearing earliest in cerebellar astrocytes and DRG cells, mirroring the regional vulnerability observed in clinical FA pathology. The severe damage in DRG non-neuronal cells predominantly SGCs- likely stems from their high mitochondrial density, which paradoxically exacerbates oxidative burden (28,37,40,44). While CNS astrocytes initially mitigate iron-mediated damage through robust buffering systems (72,78,110,112,113), they eventually succumb to cumulative stress. The elevated basal levels of frataxin observed in WT cerebrum astrocytes further suggest a cell-type-specific requirement for frataxin to sustain these defenses. However, their reduced supercomplex stability (73) creates a chronic redox imbalance, thereby lowering the threshold for metabolic failure upon frataxin depletion. Ultimately, in the DRG, the failure of supportive glia emerges as the initiating pathogenic event. This framework explains how cell-autonomous frataxin deficiency in glia precipitates the secondary, selective degeneration of sensory neurons—the defining hallmark of FA.

The progressive failure of glial defenses creates a permissive environment for ferroptosis. While previous research identified ferroptotic markers in bulk tissues (22,85), our findings reveal that signaling is strictly dependent on both cell type and anatomical region. Specifically, a ferroptosis-prone signature—increased TFR1 (iron uptake) and reduced FTH1 (iron storage)—is selectively observed in FXN^I151F^ cerebellar astrocytes and DRG cell populations as early as 21 weeks of age. This imbalance indicates a chronic failure to adequately sequester ferrous iron, expanding the labile iron pool and enhancing Fenton chemistry. In contrast, central neurons maintain a protective profile (low TFR1, high FTH1), likely through preserved iron regulatory protein 1 (IRP1) activity, allowing them to limit iron influx and delay cell death (114,115). Importantly, bulk-tissue measurements often obscure these glial defects. In our previous work using whole cerebellum, we observed increased FTH1 and decreased IRP1 (85); however, the present data indicate that this profile is largely driven by neuronal contributions, thereby masking the profound iron-handling dysfunction within astrocytes. This is consistent with evidence that susceptibility to ferroptosis varies across cell types according to their iron metabolism and antioxidant capacity (116,117). Under physiological conditions, astrocytes protect neurons by buffering iron (78); however, frataxin deficiency compromises this function, converting astrocytes into sources of neurotoxicity. Similar observations in Parkinson’s and Alzheimer’s diseases support a “glia-first” model, in which astrocytic failure drives secondary neuronal degeneration (118,119). The DRG present a more severe pathological scenario, with both neurons and non-neuronal cells exhibiting early ferroptotic signatures. Unlike the CNS, the DRG lacks a dense astrocytic network; while SGCs unsheathe neuronal somata, their protective capacities are limited (46). As a result, DRG neurons operate in a relative unprotected environment in which mitochondrial impairment caused by frataxin deficiency, precipitates ferroptosis in both cellular populations, potentially exacerbated by impaired mitochondrial transfer from SGCs (120). This contrasts with the CNS, where neurodegeneration follows a glia-driven cascade, a distinction further supported by findings in the YG8-800 model (80).

Concomitant with iron accumulation, the core antioxidant defense system collapses, thereby fulfilling the two defining requirements for ferroptosis: the presence of redox–active iron and an impaired capacity for lipid peroxidation repair. To assess the functional consequences of this convergence, we quantified lipid peroxidation—the hallmark of ferroptotic injury—across CNS and PNS tissues. In the CNS, lipid damage followed a delayed trajectory. At 21 weeks, despite evidence of iron-related stress in FXN^I151F^ mice, lipid peroxidation remained at WT levels. A significant increase in the oxidized-to-reduced BODIPY-C11 ratio emerged only at 39 weeks and was notably restricted to astrocytes. These findings identify astrocytes as the primary site of lipid membrane failure in the brain of the FXN^I151F^ mice, while neurons remain protected. In contrast, the PNS exhibited an early, widespread phenotype; by 21 weeks, both neuronal and non-neuronal cells in the DRG showed a marked increase in lipid peroxidation, suggesting a lack of regional resilience (22). To define the molecular basis of this damage, we examined GPX4, the central enzyme for detoxifying lipid peroxides. Astrocytes displayed intrinsically lower basal GPX4 levels than neurons, likely reflecting their specialization as “antioxidant factories” that prioritizes GSH export to sustain neuronal defenses (121,122), leaving them with narrower safety margins. In the CNS, GPX4 was significantly reduced in astrocytes by 21 weeks, establishing a “priming phase” in which compensatory mechanisms temporarily prevent overt lipid membrane failure until 39 weeks. In contrast, the DRG exhibited a concurrent reduction in GPX4 and a rise in lipid peroxidation by 21 weeks (22).

Given the central role of NRF2 in coordinating ferroptosis resistance (77,123), we investigated its contribution to cellular defense. In WT animals, astrocytes exhibit significantly higher basal NRF2 levels than neurons, stabilized via p35/Cdk5 signaling (94). This allows astrocytes to serve as the primary source of GSH precursors for neuronal protection (30). This function is linked to astrocytic mitochondrial architecture; unlike neurons, astrocytes maintain a larger pool of free complex I, generating basal ROS that serve as essential signaling cues for NRF2 readiness (73). In the FXN^I151F^ model, this homeostatic advantage erodes. While CNS neurons showed a transient increase in NRF2 at 21 weeks, astrocytes exhibited a marked decline by 39 weeks. This suggest that frataxin deficiency converts mitochondrial ROS from beneficial signals into a toxic burden that eventually exhausts the NRF2 pathway. This collapse is particularly evident in the cerebellum, where astrocyte dysfunction drives neuronal vulnerability (84). By 39 weeks, cerebellar astrocytes showed a sharp decline in NRF2 coinciding with lipid peroxidation, whereas neurons exhibited sustained NRF2 reduction without immediate lipid damage, implying reliance on alternative protective mechanisms. In the DRG, the phenotype was more severe: both neurons and non-neuronal cells exhibited early NRF2 loss by 21 weeks, consistent with reported defects in nuclear localization (22). This NRF2 depletion represents a “double hit,” simultaneously compromising intrinsic neuronal defenses and the essential glial metabolic support required for survival. Ultimately, the failure of the NRF2 system in both the CNS and PNS underscores the transition from a compensated oxidative state to an unrecoverable ferroptotic environment.

The profound molecular stress within the glial compartment is reflected in the development of astrogliosis, marked by increased GFAP expression (95). The reduction of NRF2 in FXN^I151F^ astrocytes likely exacerbates oxidative stress, promoting this reactive phenotype. Primary redox imbalances in astrocytes are known to induce proinflammatory states that accelerate neurodegeneration (124). Accordingly, we observed early and sustained GFAP increases in the cerebrum, cerebellum, and DRG, consistent with robust glial reactivity in FA models (80). This widespread response suggests that compromised NRF2-mediated defenses drive a reactive glial state that contributes to disease progression.

In summary, this study supports a paradigm shift in FA pathology by identifying glial cells as the earliest and most profoundly affected compartment (Fig. 9). Disease progression follows a hierarchical sequence: mitochondrial dysfunction triggers iron overload, leading to the collapse of the NRF2–GPX4 axis, ferroptosis, and reactive gliosis. This framework explains regional vulnerability and clarifies why the DRG is the primary site of pathology. In the CNS, severe defects like supercomplex disruption and ferroptosis remain largely confined to astrocytes. Their robust antioxidant capacity provides a transient buffer that delays neuronal degeneration. Conversely, DRG neurons rely on less resilient SGCs, which offer limited antioxidant and iron-handling support. This heightened vulnerability results from the lower baseline capacity of SGCs and their extreme physical proximity to the neuronal soma, leading to early, concurrent failure in both populations. This aligns with findings where regional susceptibility is dictated by glial antioxidant capacity (124) and recent work showing that frataxin deficiency in proprioceptors alters SGC and macrophage gene transcription (125). These distinctions carry significant therapeutic implications. Interventions must prioritize preserving glial function to halt neurodegenerative cascades, as glia are active drivers of oxidative stress and neuroinflammation (126). In this context, NRF2 induction emerges as a promising strategy. Compounds such as omaveloxolone, leriglitazone, and vatiquinone have shown efficacy by modulating NRF2-related pathways (22,93,127,128), with the approval of omaveloxolone further validating NRF2 stabilization (129). Reinforcing astrocytic NRF2 may preserve the “glial shield” and interrupt the ferroptotic cascade before it reaches vulnerable neurons (105).

**Figure 9.**
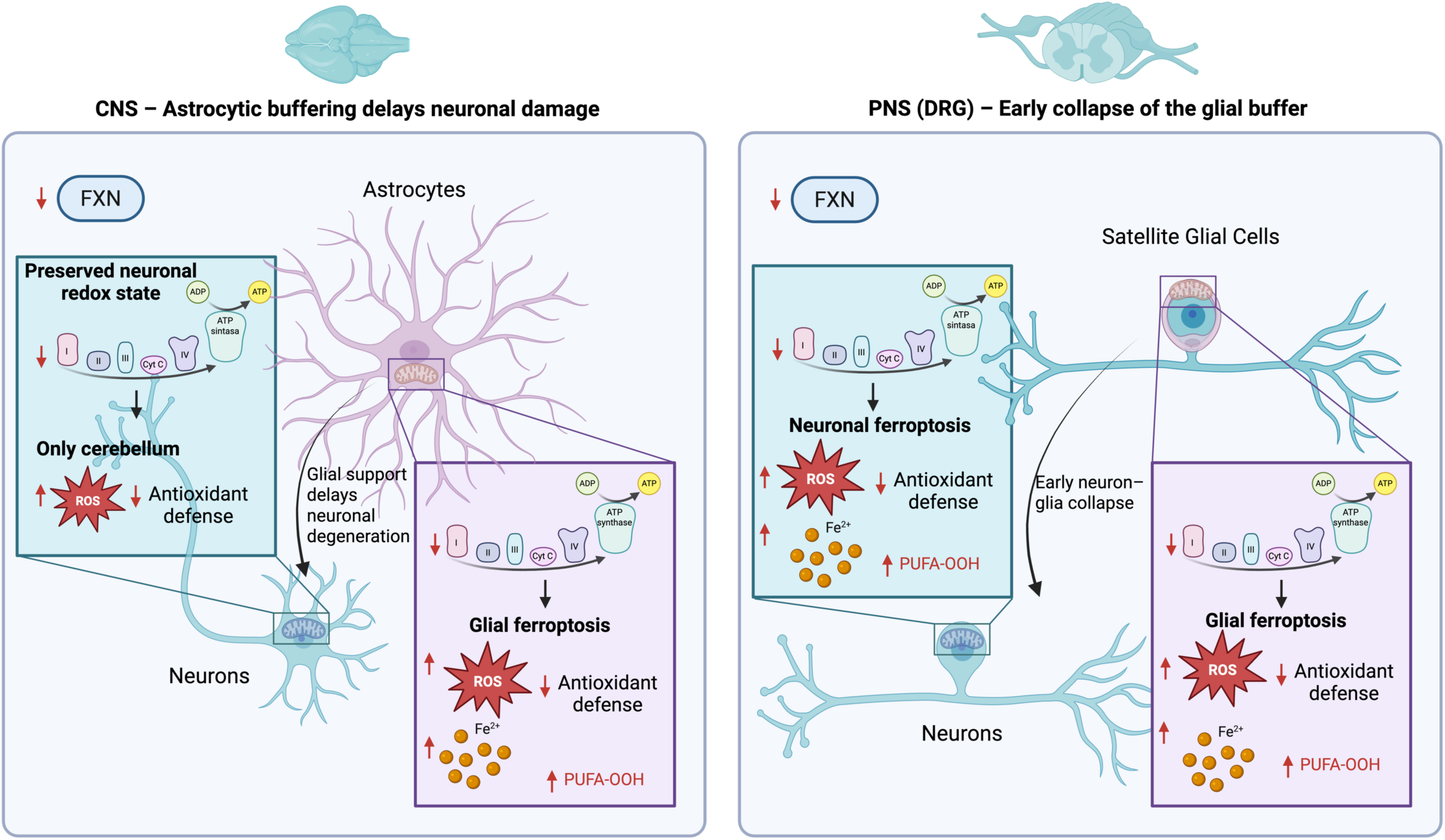
Model of differential vulnerability to frataxin deficiency in the CNS and PNS. Schematic illustration of the glial buffer hypothesis in FA. CNS – Astrocytic buffering delays neuronal degeneration (left): Frataxin deficiency induces mitochondrial dysfunction, including impaired respiratory chain activity and ATP production. In the CNS, astrocytes are initially more affected but possess strong antioxidant defenses that enable them to buffer iron and ROS, thereby preserving neuronal redox balance. This astrocytic support delays lipid peroxidation (PUFA-OOH) and ferroptotic death in neurons, with neuronal vulnerability largely restricted to the cerebellum. PNS (DRG) – Early collapse of the glial buffer (right): In the DRG, SGCs exhibit limited antioxidant reserves. Frataxin deficiency causes an early breakdown of these defenses, leading to iron accumulation, increased ROS, and elevated lipid peroxidation. The close tight anatomical association between SGCs and neuronal cell bodies promotes a rapid, concurrent collapse of glial-neuronal homeostasis, triggering early and widespread ferroptosis in both cell types.

## Conclusion

This study provides the first systematic, cell-type comparison of neurons and glial cells across CNS and PNS regions in FA, revealing that glial dysfunction—rather than neuronal failure—is the primary driver of regional degeneration. By mechanistically linking mitochondrial impairment, iron dysregulation, and ferroptosis to regional differences in glial support capacity, these findings underscore the preservation of glial homeostasis as a critical therapeutic priority and position non-neuronal compartments as central targets for disease-modifying interventions.

## Data availability

The data generated and analyzed during the current study are available in the figures and supplementary materials, and have been deposited in a citable, publicly accessible repository (Mendeley data: https://data.mendeley.com/datasets/bghbdxszxc/2).

## Ethics statement

All animal procedures were conducted in accordance with the National Guidelines for the Regulation of the Use of Experimental Laboratory Animals from the Generalitat de Catalunya and the Government of Spain (Article 33.a, 214/1997) and approved by the University of Lleida Experimental Animal Ethics Committee (CEEA).

## Acknowledgments

We gratefully acknowledge Roser Pané for outstanding technical support, and Anaïs Panosa from the Microscopy and Flow Cytometry facilities for assistance with confocal imaging and flow cytometry. We also thank the Cell Culture facility at UdL, as well as the Neuroenergetics and Metabolism Group facilities (IBFG, USAL-CSIC), for their support in conducting experiments. Additionally, we are grateful to Ester Vilapriñó for assistance with statistical analyses and to Alaó Gatius for help with the quantification of confocal images.

## Funding

This work was supported by: 1) Grant PID2020–118296RB-I00 funded by MICIU/AEI /10.13039/501100011033; 2) Grant PDC2021–120758-I00 funded by MICIU/AEI /10.13039/501100011033 and by the European Union NextGenerationEU/ PRTR; 3) Grant PID2023–148128OB-I00 funded by MICIU/AEI /10.13039/501100011033 and by FEDER, EU; 4) Project 2021-SGR 00323 funded by Generalitat de Catalunya. A.S-A received first a Ph.D. fellowship from the Generalitat de Catalunya and after, she held predoctoral fellowship “Ajuts al Personal Investigador en Formació “ from IRBLleida/Diputació de Lleida. M.P-C received a PhD fellowship from the Generalitat de Catalunya and after, and after she held predoctoral fellowship “Beca de Suport per la finalització de la Tesis Doctoral” from IRBLleida/Diputació de Lleida. M.P-G received first a PhD fellowship from the Universitat de Lleida and after, and after she held predoctoral fellowship “Beca de Suport per la finalització de la Tesis Doctoral” from IRBLleida/Diputació de Lleida J.P.B. is funded by MICIU/AEI (PID2022-138813OB-I00 /10.13039/501100011033 and FEDER, UE), la Caixa Foundation (grant agreement LCF/PR/HR23/52430016), the European Union’s Horizon Europe research and innovation program under the MSCA Doctoral Networks 2021 (ref. 101072759); FuEl ThE bRaiN In healtThY aging and age-related diseases, ETERNITY, and the European Research Council (ERC) Advanced Grant NeuroSTARS (ref. 101199747).

## Declaration of competing interest

The authors declare that they have no known competing financial interests or personal relationships that could have appeared to influence the work reported in this paper.

## Author contributions

Conceptualization, data curation, methodology, investigation and visualization: A.S-A., M.P-C., I.M-R., M.P-G., F.D., J.T., J.P.B., J.R. and E.C. Formal analysis and validation: A.S-A., J.R. and E.C. Resources, funding acquisition and supervision: J.T., J.P.B., J.R. and E.C. Sofware, and writing original draft: A.S-A., and E.C. Writing – review and editing: A.S-A., I.M-R., J.T., J.P.B., J.R. and E.C. All authors have read and agreed to the published version of the manuscript.

## Abbreviations

BNGE: Blue native gel electrophoresis
BSA: bovine serum albumin
CNS: central nervous system
CBB: Coomassie Brilliant Blue
DMEM: Dulbecco’s modified Eagle’s medium
DPBS: Dulbecco’s phosphate-buffered saline
DRG: dorsal root ganglia
ECAR: extracellular acidification rate
EBSS: Earle’s Balanced Salt Solution
FA: Friedreich ataxia
FCCP: carbonyl cyanide-p-trifluoromethoxyphenylhydrazone
FTH1: ferritin heavy chain GFAP glial fibrillary acidic protein
GPX4: glutathione peroxidase 4
GS: glutamine synthetase
GSH: glutathione
IRP1: iron regulatory protein 1
MAP2: Microtubule-Associated Protein 2
NRF2: nuclear factor erythroid 2-related factor 2
OCR: oxygen consumption rate
OXPHOS: oxidative phosphorylation
PBS: phosphate-buffered saline
PFKFB3: 6-phosphofructo-2-kinase/fructose-2 6-bisphosphatase 3
PNS: peripheral nervous system
PUFA: polyunsaturated fatty acid
ROS: reactive oxygen species
SDS-PAGE: SDS-polyacrylamide gel electrophoresis
SGC: satellite glial cell
TFR1: transferrin receptor 1.

## Competing interests

### Declaration of interest

The authors declare no competing interests.

## Supplementary material

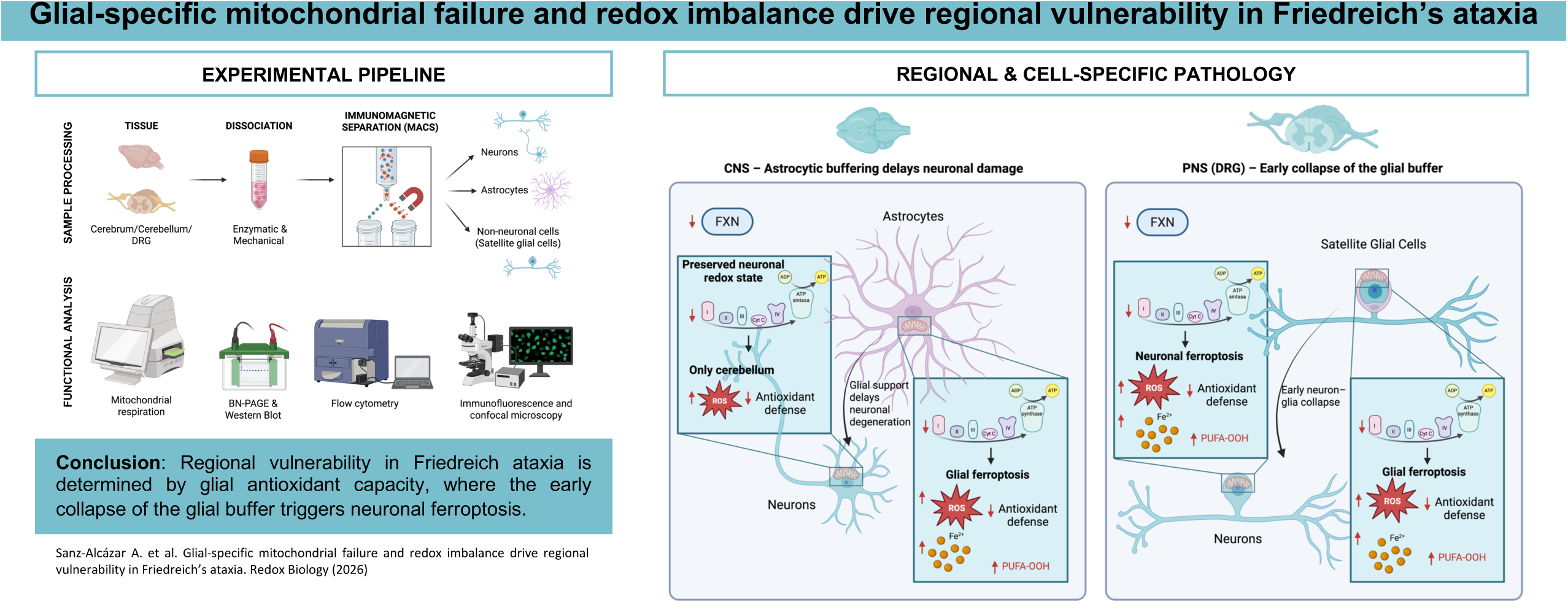

## Notes

https://data.mendeley.com/datasets/bghbdxszxc/2

### Competing Interest Statement

The authors have declared no competing interest.

